# π-MNovo improves de novo peptide sequencing through microbial-domain adaptation and evidence-guided candidate selection

**DOI:** 10.64898/2026.09.22.753549

**Authors:** Jingwen Ye, Xin Zhang, Boyan Sun, Tianze Ling, Zhendong Liang, Jingyi Dong, Yuan Zheng, Hui Jin, Liming Jin, Zikai Hao, Leyuan Li, Cheng Chang

## Abstract

The high taxonomic and strain-level diversity of microbial communities makes it difficult for reference databases to fully represent the protein sequences present in metaproteomic samples, limiting database-dependent peptide identification. De novo peptide sequencing can recover peptide sequences directly from tandem mass spectra without relying on reference databases, providing complementary peptide evidence for metaproteomics. However, most existing de novo sequencing models were developed largely from non-microbial proteomic data and lack specific adaptation to microbial spectra. Here, we established a dedicated microbial spectral resource comprising more than 10 million annotated high-quality tandem mass spectra from 72 cultured microbial isolates, with peptide digests from each isolate separated into five high-pH reversed-phase fractions to increase the opportunity for deeper and more diverse peptide sampling. Using this resource, we developed π-MNovo, which combines adaptation to microbial spectra with evidence-guided candidate selection to improve full-length peptide sequencing precision. On a seven-species external benchmark, π-MNovo achieved 66.30% complete-peptide recall under the same residue-mass-based complete-peptide matching criterion applied to all models, exceeding four published models by 6.9–21.7% in relative recall, with consistent gains across species and peptide-length groups. In two independent metaproteomic datasets, π-MNovo increased reference-supported taxonomic coverage, while synthetic-community analysis recapitulated the designed abundance ranking. From pFind-unidentified spectra, π-MNovo recovered 1,587 pFind-unreported, reference-matched peptides that passed spectrum-evidence filtering. These results show that π-MNovo expands recoverable peptide evidence and enhances the utility of de novo sequencing for metaproteomic analysis.

## Introduction

Metaproteomics connects microbial community composition with functional activity by identifying and quantifying community-derived proteins. Interpreting these measurements requires assigning MS/MS spectra to peptide sequences, which provide evidence for protein identification and taxonomic annotation. Conventional database searching assigns spectra to sequences in a reference protein database, making database sequence coverage and accuracy central to the analysis. In complex microbial communities, both sample-specific metagenomic databases and public reference databases may incompletely represent the peptide sequences present in the sample. Sample-specific metagenomes can be limited by insufficient sequencing depth and incomplete assembly, whereas public databases may omit species without sequenced genomes and contain reference strains that differ from the strains actually present. Consequently, some sample-derived peptides may have no exact match in the database used for searching^1^. Recovering sequence evidence beyond the coverage of a conventional search is therefore important for improving sequence recovery in metaproteomic analysis.

*De novo* peptide sequencing addresses this limitation by inferring sequences directly from MS/MS spectra. Its appl<u>icability</u> for metaproteomics has already been demonstrated: MetaNovo uses *de novo* sequence tags to construct sample-adapted search databases^2^, and novoMP combines *de novo* sequencing with multilayer quality filtering and FDR validation to expand sequence databases and taxonomic coverage^3^. Advances in deep learning have also improved the underlying sequencing models. Casanovo uses Transformer-based spectrum-to-peptide translation^4^; π-HelixNovo introduces complementary-spectrum encoding to enhance fragment-ion information^5^; π-HelixNovo2 further integrates complementary-spectrum encoding with bidirectional decoding and peptide-level quality control^6^; and π-PrimeNovo combines a non-autoregressive Transformer with connectionist temporal classification and precursor-mass-constrained decoding^7^. These developments provide a foundation for recovering peptide evidence from microbial spectra.

Applying these models to microbial samples nevertheless raises a cross-domain generalization challenge. Most existing models have been pretrained predominantly on spectra from human proteome spectral libraries or synthetic peptides based on human protein sequences^8^, whereas metaproteomics encompasses highly diverse microbial proteomes; for example, an individual gut microbiome may contain 200–500 species. Previous evaluations and fine-tuning studies have established the feasibility of transferring existing models to microbial spectra, but improving complete-peptide recovery in microbial proteomes remains an important practical challenge. Effective domain adaptation requires not only a large number of annotated spectra but also broad coverage of the microbial peptide sequence space on which the model must generalize. In single-shot data-dependent LC–MS/MS analyses, finite precursor-selection capacity favors abundant ions and leaves many co-eluting peptides unsampled. Offline high-pH reversed-phase fractionation distributes peptide complexity across multiple runs, increasing the opportunity to acquire a more diverse set of annotated peptide-spectrum matches (PSMs). This within-isolate peptide depth complements the taxonomic breadth provided by sampling multiple microbial isolates. Long peptides are particularly demanding because residue-level errors and missing fragment evidence can prevent recovery of the complete sequence^9^. At the same time, longer sequences can provide more specific reference-based taxonomic evidence^10–11^. Improving microbial peptide sequencing therefore requires attention to both overall recovery and the specific challenges of long peptides.

Domain adaptation can improve the way candidates are generated, but generation alone does not determine which sequence is ultimately reported. A reference-matching candidate may already be present in the decoded candidate pool yet rank below an incorrect alternative. Precursor-mass consistency, peptide length and fragment-ion support offer complementary information for resolving such cases. However, the value of each evidence source varies across spectra, so using an additional source indiscriminately may also displace correct predictions. This motivates a candidate-selection strategy that combines different evidence types while retaining the default ranking when an alternative is insufficiently supported.

Here we asked whether microbial-domain adaptation and better use of candidate-level evidence could jointly improve complete-peptide recovery from microbial MS/MS spectra. To provide both taxonomic breadth and within-isolate peptide depth, we first constructed a microbial spectral resource comprising more than 10 million annotated PSMs from 72 extensively fractionated isolate-derived samples, with development and internal test sets partitioned by canonical peptide sequence. Using this resource, we developed π-MNovo by adapting an existing non-autoregressive backbone and combining shared candidate generation with evidence-guided ranking and conservative expert routing. We assessed predictive performance using spectra from external microbial species, then used real metaproteomes and a defined synthetic community to examine whether improved peptide recovery translated into broader reference-supported evidence for taxonomic and community analyses.

## Results

### A fractionated, isolate-derived microbial spectral resource with peptide-disjoint partitions

π-MNovo performs *de novo* peptide sequencing directly from MS/MS spectra, providing sequence evidence for subsequent protein mapping and taxonomic interpretation (Fig. 1a). To support microbial-domain adaptation, we constructed an annotated spectral resource designed to combine taxonomic breadth across microbial isolates with broader peptide sampling within each isolate (Supplementary Data 1).

**Fig. 1.**
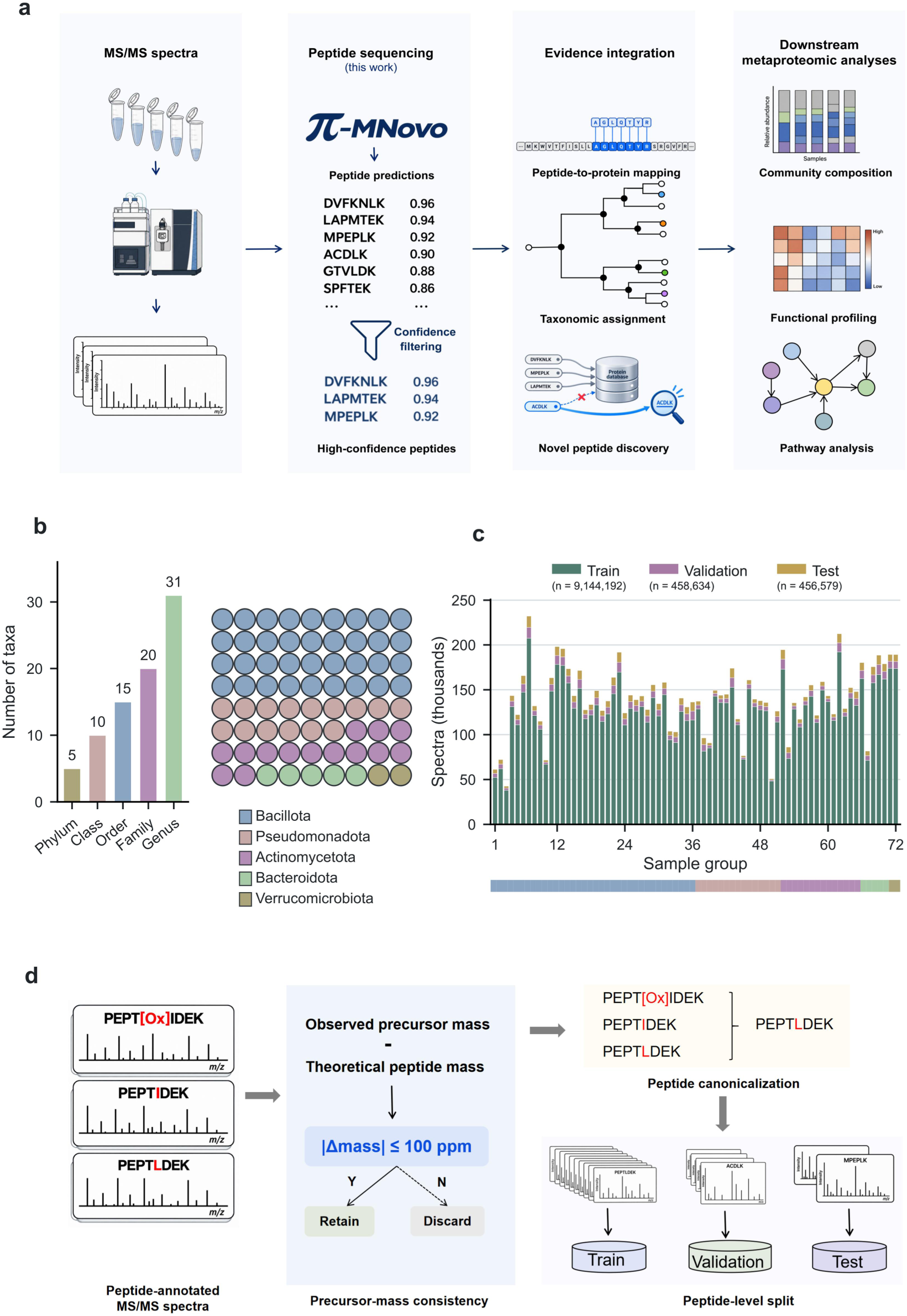
Construction of the microbial peptide-spectrum resource. a, Positioning of π-MNovo within the metaproteomic analysis workflow. π-MNovo converts MS/MS spectra into high-confidence peptide predictions, which provide sequence evidence for protein mapping, taxonomic assignment and novel-peptide discovery, followed by downstream community, functional and pathway analyses. b, Taxonomic profiles of the 72 cultured bacterial isolates included in the resource. The bars show the number of distinct taxa at each rank; each circle represents one retained isolate, colored by phylum. c, Retained spectra in thousands for each sample group, stacked by training, validation and test partition. Isolate-level sample groups comprising the available fraction files are ordered by phylum, as indicated by the color strip below the bars. Counts reflect the final resource after filtering and per-key capping. d, Precursor-mass consistency filtering, peptide canonicalization and canonical-peptide-level partitioning. Peptide-annotated spectra with a precursor-mass error of ≤ 100 ppm were retained; modification annotations were removed and I/L-equivalent sequences were assigned to mutually exclusive training, validation and test partitions.

The final collection comprised 72 cultured gut bacterial isolates recovered from healthy human donors. Only isolates for which the peptide-based Unipept assignment was concordant with the corresponding 16S rRNA gene assignment, and for which the 16S rRNA gene sequencing results were consistent with a pure culture, were retained. For each retained isolate, the consensus taxonomic assignment was mapped to the corresponding NCBI Taxonomy identifier (TaxID) and used to retrieve the relevant proteome(s) from UniProtKB, thereby generating an isolate-specific protein sequence database for second-pass pFind searching. The 72 retained isolates spanned 5 phyla, 10 classes, 15 orders, 20 families and 31 genera (Fig. 1b; Supplementary Data 2). Bacillota was the most represented phylum (n = 36, 50%), followed by Pseudomonadota (n = 15, 21%), Actinomycetota (n = 14, 19%), Bacteroidota (n = 5, 7%) and Verrucomicrobiota (n = 2, 3%; Fig. 1b).

To increase the opportunity for broader peptide sampling within each isolate, each isolate digest was subjected to five-fraction high-pH reversed-phase prefractionation before LC–MS/MS analysis. The experimental design comprised 360 planned fraction-level analyses. Deep proteomic profiling was designed to generate five fractions per isolate (Supplementary Fig. S1). Fourteen fraction samples from 12 isolates were accidentally lost during experimental processing. Together with ten files contributed by one isolate, this resulted in 351 MGF files from 72 isolates being available for resource construction. After quality control and per-peptide spectrum capping, the final resource contained 10,059,405 annotated spectra. Spectra were partitioned according to canonical peptide sequence into training, validation and internal test sets containing 9,144,192 (training), 458,634 (validation) and 456,579 (internal) spectra, respectively (Fig. 1c). Before partitioning, peptide-annotated spectra were filtered for precursor-mass consistency and canonicalized by removing modification annotations and treating I and L as equivalent (Fig. 1d). Canonical peptide keys were mutually exclusive across the three primary partitions, such that repeated spectra and modified forms derived from the same underlying peptide sequence remained within a single partition. Spectrum counts were capped at 120 per canonical peptide key in the training partition and at five per key in each of the validation and test partitions. Search settings and partitioning procedures are detailed in Supplementary Tables S1 and S2 and Supplementary Methods 1–2.

Long peptides were also well represented in the resource. Peptides of at least 20 residues accounted for 2.13 million spectra, corresponding to 21.22% of all retained spectra (Supplementary Fig. S2). Within the training partition, a fixed subset of one million spectra was used for backbone adaptation, whereas a peptide-disjoint set of 996,518 spectra was used to develop the candidate rankers, length predictor, expert paths and router. Together, the resource provided a large peptide-disjoint basis for model development and internal evaluation while retaining broad taxonomic representation and substantial coverage of long peptides.

### π-MNovo improves complete-peptide recall across external microbial species

Using this resource, we developed π-MNovo around a shared non-autoregressive backbone. CTC beam search and precursor-mass-constrained decoding use the output of a single backbone forward pass to generate a shared candidate pool (Fig. 2a). The default mass-aware ranker and two specialized ranking paths rescore the same candidate pool; the specialized paths incorporate peptide-length and fragment-ion evidence. A conservative router selects an expert path only when the router’s preference for that path over the default exceeds a fixed margin; otherwise, the default prediction is retained. The selected sequence is returned with a calibrated Score. This architecture separates candidate generation from evidence-guided selection, allowing additional evidence to refine the final choice without generating a new pool for each path. Architecture and component-development procedures are detailed in Supplementary Methods 3–7, and the final component configuration is provided in Supplementary Data 3.

**Fig. 2.**
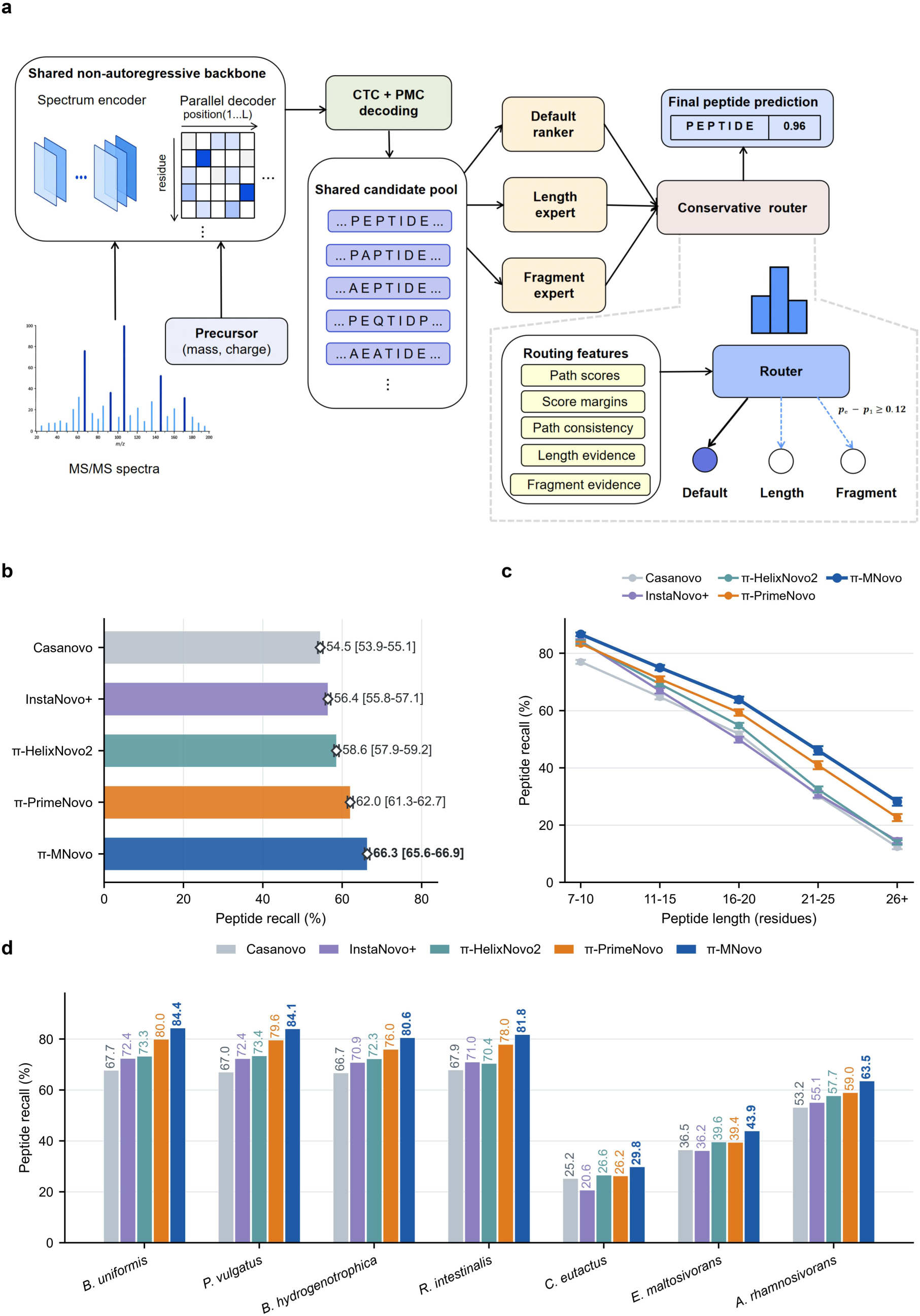
Architecture and complete-peptide recall on the external microbial benchmark. a, The shared non-autoregressive backbone supplies CTC beam-search and precursor-mass-constrained (PMC) candidates to three ranking paths: the default ranker (R1), length expert (R2) and fragment expert (R3). The conservative router switches from R1 when pₑ − p₁ ≥ 0.12, where pₑ is the larger expert-path softmax probability and p₁ is the R1 probability. The selected candidate’s R1 score is mapped to the final sequence Score by Platt scaling. b, Micro-averaged recall across 203,204 spectra from seven species. Bars and terminal diamonds denote recall. Error bars show 95% percentile intervals from 5,000 paired hierarchical bootstrap replicates, with species treated as fixed strata, runs resampled within species and spectra resampled within runs. c, Recall by reference-peptide length, using the same bootstrap procedure. The five length groups contain 39,745, 68,309, 49,004, 26,724 and 19,365 spectra; 57 spectra shorter than seven residues contribute to the overall analysis. d, Recall for each species and model. Model colors are consistent across b–d. Complete-peptide matching follows the mass-aware rule in Methods; empty and invalid predictions remain in the recall denominator.

We evaluated π-MNovo on an external benchmark of 203,204 spectra from seven microbial species, drawn from PXD037923^12^, PXD012448^13^, PXD014174^14^, PXD028575^15^ and PXD021084^16^. The species were absent from backbone training, and reference peptides sharing a canonical key with the backbone-training set were excluded. Each species contributed at most 30,000 spectra. We compared π-MNovo with the released implementations of Casanovo^4^, InstaNovo+^8^, π-HelixNovo2^6^ and π-PrimeNovo^7^ using the same spectra and mass-aware complete-peptide matching rule (Supplementary Tables S3 and S4; Supplementary Method 8; Supplementary Data 4 and 5).

On this benchmark, π-MNovo achieved 66.30% complete-peptide recall. This exceeded the four comparison models by 4.27–11.81 percentage points, corresponding to 8,671–24,002 additional complete predictions matching the benchmark reference (Fig. 2b). π-MNovo ranked first in all seven species (Fig. 2d). Recall declined with peptide length for all models. The margin over the highest-recall comparator in each length group increased from 2.37 percentage points for 7–10-residue peptides to 5.55 points for peptides of at least 26 residues. The gains for the 11–15-, 16–20- and 21–25-residue groups were 3.97, 4.49 and 5.17 points, respectively (Fig. 2c). π-MNovo achieved 28.17% recall in the longest group. Thus, π-MNovo improved complete-peptide recall across species and showed its largest absolute gains for longer peptides.

### Sequential component additions improve complete-peptide recall

The external benchmark established the overall performance gain; we used the internal ablation set to quantify changes associated with sequential component additions and to compare decoding beam widths. Across 456,579 spectra, complete-peptide recall rose from 61.46% for the base model to 63.32% after microbial-domain adaptation, 63.73% after weight interpolation, 65.88% after mass-aware reranking and 66.02% after conservative routing (Fig. 3a). The largest sequential increments followed microbial-domain adaptation and default reranking (Supplementary Data 6).

**Fig. 3.**
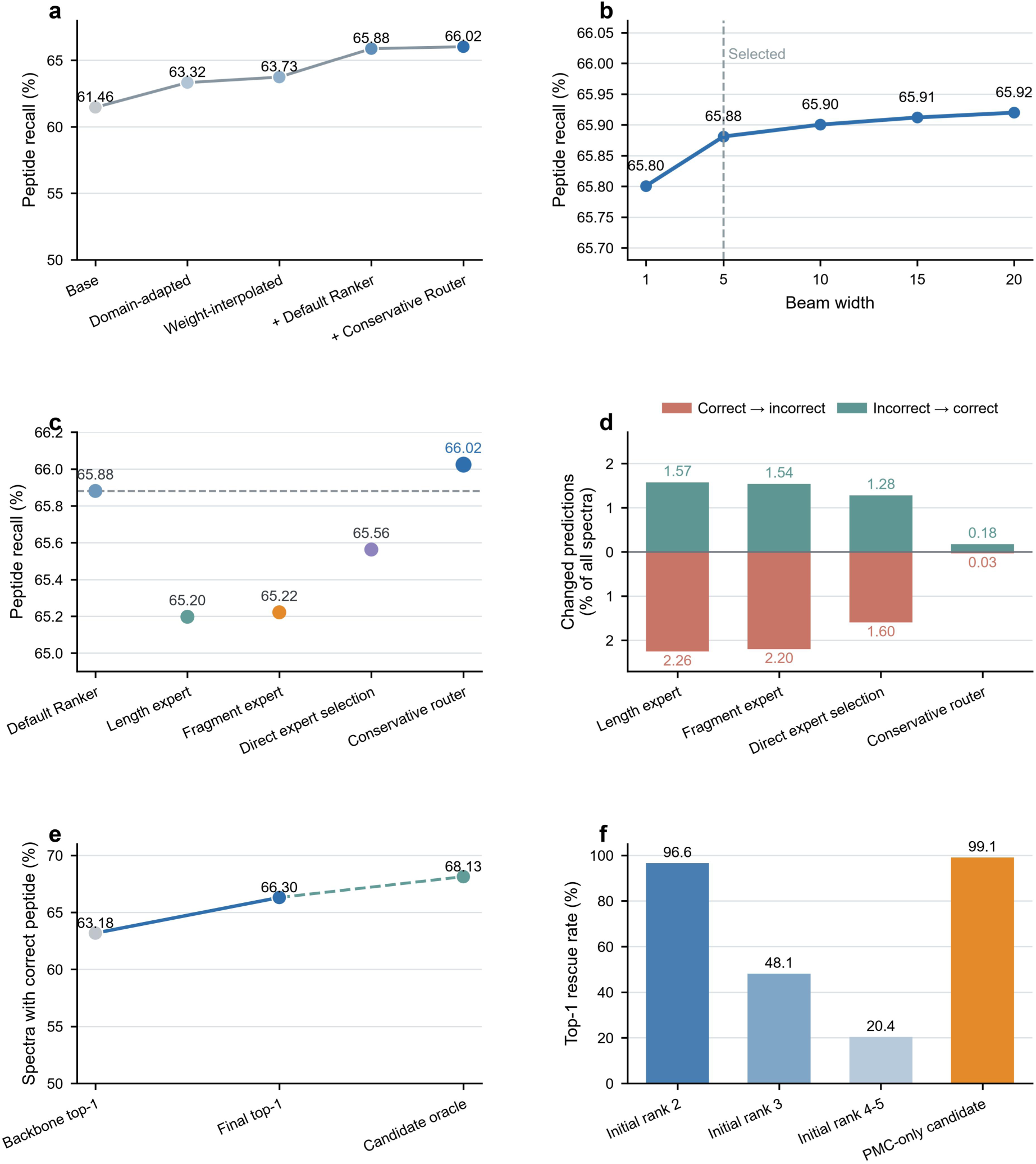
Sequential component analysis, candidate width and expert routing. Panels a–d use the internal ablation set comprising 456,579 spectra; e and f use the 203,204-spectrum external seven-species benchmark. a, Recall after successive additions of domain adaptation, weight interpolation, the default ranker R1 and conservative routing. b, Recall at different CTC beam widths with the backbone, R1 weights and PMC candidates held fixed. “Selected” and the dashed line indicate the beam width of 5 used in the released model. c, Recall when each displayed selection strategy is applied to all spectra. The horizontal line marks R1 recall; length and fragment experts add their respective evidence to the R1 score. d, Changes relative to R1: “Correct→ incorrect” indicates harmful switches and “Incorrect → correct” indicates beneficial switches. Percentages use all internal ablation spectra as the denominator. e, “Backbone top-1” is the proportion of spectra whose initially top-ranked candidate matches the reference; “Final top-1” is the proportion whose final selected candidate matches; “Candidate oracle” is the proportion with at least one matching candidate anywhere in the shared pool. f, Selection rates for spectra whose shared candidate pool contains a reference-matching sequence that is absent from the initial top-1 output, grouped by the sequence’s initial rank or exclusive origin from PMC. Each rate uses the corresponding subgroup of spectra with a reference-matching candidate in the pool as its denominator.

Increasing candidate-search width produced only a small change in this configuration. With the backbone and reranking weights fixed, increasing the CTC beam width from 5 to 20 changed recall by 0.0388 percentage points, corresponding to 177 additional correct predictions (Fig. 3b). The released configuration uses a beam width of 5.

Selective routing outperformed applying an expert path to every spectrum. Applying the length expert, the fragment expert or direct score-based path selection to every spectrum did not outperform the default reranker (Fig. 3c). Conservative routing, by contrast, corrected predictions for 809 spectra but changed 152 initially correct predictions to incorrect ones, yielding a net gain of 657 correct predictions (Fig. 3d). The default path supplied 98.90% of outputs, indicating that the gain from routing came from a small fraction of predictions. These results highlighted the distinction between candidate generation and candidate selection. On the external seven-species benchmark, the initially top-ranked candidate matched the reference for 63.18% of spectra, whereas a matching sequence was present in the shared candidate pool for 68.13%. Evidence-guided selection raised final recall to 66.30%, closing much of this gap without regenerating candidates (Fig. 3e). Recovery varied by initial candidate rank and origin. For spectra in the initial rank-2, rank-3 and rank-4–5 groups, final predictions matched the reference in 96.6%, 48.1% and 20.4% of cases, respectively. The selection rate for candidates supplied only by the PMC branch was 99.1% (Fig. 3f). Across the benchmark, candidate selection corrected predictions for 6,376 spectra and changed 28 initially correct predictions to incorrect ones, yielding a net gain of 6,348 correct predictions. Thus, candidate selection could correct some initial prediction errors. After final selection, recall remained 1.83 percentage points below the fixed-pool ceiling.

### π-MNovo expands peptide and taxonomic evidence in human metaproteomes

We next evaluated model performance on real metaproteomic spectra, which differ from isolate spectra in sample complexity and biological composition. The evaluation included the fecal metaproteomic dataset PXD020786^17^ (151,238 spectra) and the salivary metaproteomic dataset PXD055269^18^ (583,531 spectra). Although absolute recall was lower in these human microbiome-derived metaproteomes, π-MNovo achieved the highest recall across 734,769 pooled reference-annotated spectra. Its recall of 28.53% exceeded the four comparison models by 2.81–14.69 percentage points (Fig. 4a). Recall was 32.62% for reference peptides present in backbone-adaptation data and 27.09% for peptides absent from those data, with corresponding gains of 3.27–16.62 and 2.65–14.01 points. The advantage therefore extended beyond sequences encountered during adaptation.

**Fig. 4.**
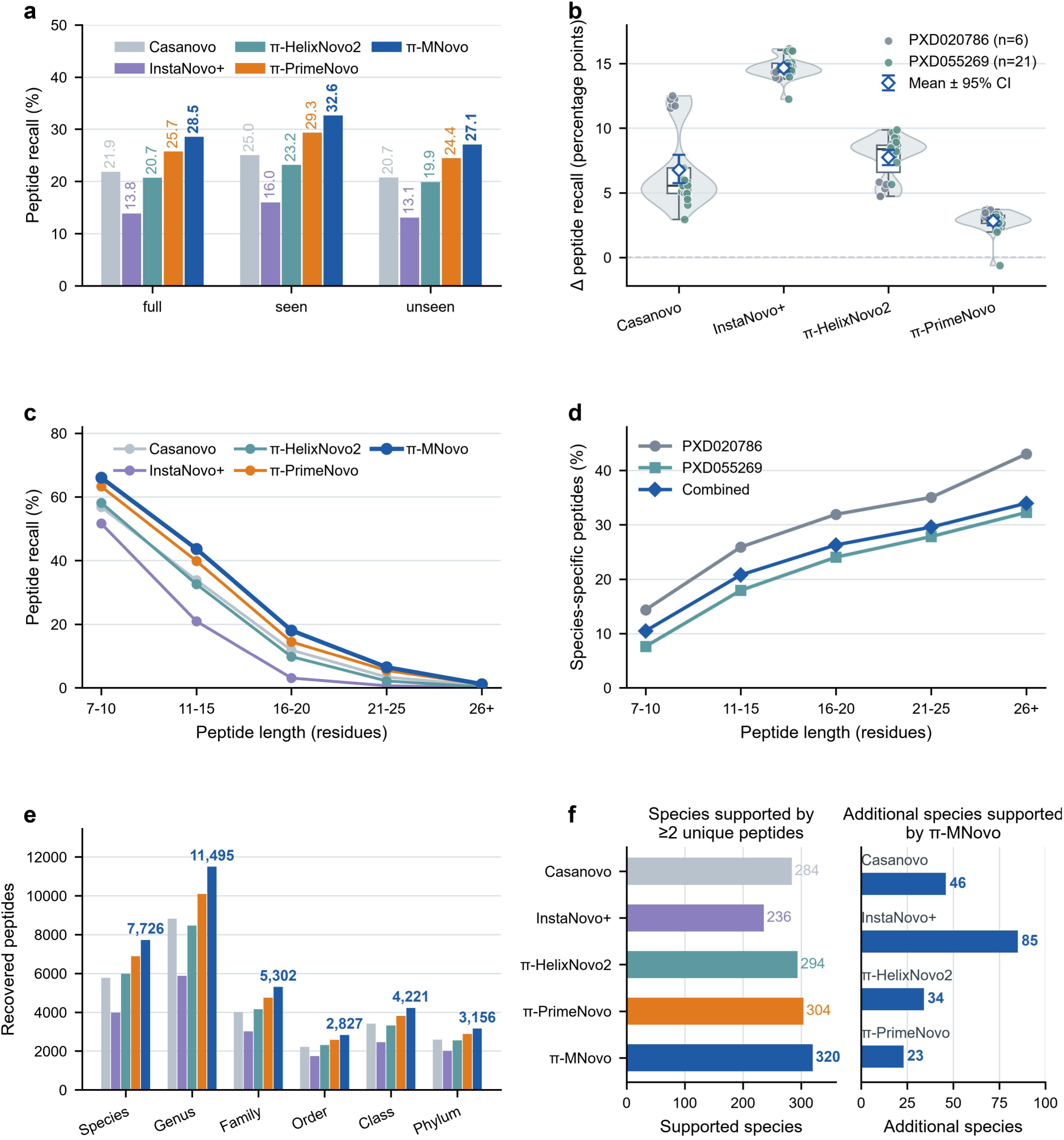
Complete-peptide recall and taxonomic evidence in real metaproteomes. a, Recall across 734,769 pooled reference-annotated spectra. “Full” includes all spectra; “seen” (n = 191,447) and “unseen” (n = 543,322) denote canonical reference peptides present in or absent from π-MNovo backbone-training data, respectively. b, Run-level recall differences, calculated as π-MNovo minus each comparator in percentage points. Gray and teal points represent six PXD020786 runs and 21 PXD055269 runs, respectively. Violins show the distributions; boxes show medians and interquartile ranges, with whiskers extending to the most extreme values within 1.5 times the interquartile range. Open diamonds show mean differences, and error bars show 95% percentile intervals from 10,000 bootstrap resamples of the 27 runs. c, Recall by reference-peptide length. d, Fraction of nonredundant reference peptides with a species-level LCA, stratified by project and peptide length. e, Nonredundant reference-matched peptides recovered by each model, counted exclusively at their assigned LCA rank. f, Species supported by at least two distinct species-LCA peptides (left) and species supported by π-MNovo but absent from each comparator’s supported set (right). Model colors are consistent across a, c, e and the left part of f. Taxonomic specificity in d–f is defined within the reference-mapping space described in Methods.

The recall advantage was observed in both projects, despite their biological and technical differences. π-MNovo achieved recall of 45.77% in PXD020786 and 24.06% in PXD055269, exceeding the comparison models by 3.51–14.25 and 2.63–14.80 percentage points, respectively. Across the 27 LC–MS/MS runs, the mean recall difference was positive for every comparator, although the magnitude varied and one PXD055269 run had a negative difference relative to π-PrimeNovo (−0.62 percentage points; Fig. 4b). π-MNovo also ranked first in every peptide-length stratum in the pooled analysis, although recall declined with increasing length for all models (Fig. 4c).

We next examined whether improved peptide recovery translated into broader taxonomic evidence, using nonredundant reference-matched peptides assigned to their lowest common ancestor (LCA; Supplementary Method 9). Longer reference peptides were more frequently assigned at the species level. In the pooled real metaproteomes, the fraction of nonredundant reference peptides assigned to a species-level LCA rose from 10.5% for 7–10-residue peptides to 34.0% for peptides of at least 26 residues, with the same trend in both projects (Fig. 4d). With each peptide counted only at its assigned LCA rank, π-MNovo recovered the most reference-matched peptides at every rank from species to phylum (Fig. 4e). It recovered 11,495 genus-LCA peptides, compared with 5,871–10,096 for the four comparison models, and 7,726 species-LCA peptides. Using a threshold of at least two distinct species-LCA peptides, π-MNovo supported 320 species, compared with 284, 236, 294 and 304 for Casanovo, InstaNovo+, π-HelixNovo2 and π-PrimeNovo, respectively. Pairwise set comparisons identified 46, 85, 34 and 23 species supported by π-MNovo but not by each comparator, whereas 10, 1, 8 and 7 were supported only by the corresponding comparator (Fig. 4f). Thus, the additional peptides increased the evidence available at individual taxonomic ranks and the number of species meeting the reference-based support criterion (Supplementary Data 7).

### π-MNovo recapitulates synthetic-community abundance and recovers additional sequence evidence from unidentified spectra

We next tested whether abundances estimated from de novo peptide intensities reproduced the designed abundance structure of the synthetic community PXD00611819. The cellular reference for the abundance-gradient mixtures U1–U4 contained 23 members: 21 bacterial species, one archaeal species and one microalgal species. π-MNovo supported all 23 members with at least two species-LCA peptides in each of the eight technical acquisitions. Abundances were estimated separately for π-MNovo and pFind from their respective PANDA20 peptide intensities. π-MNovo reproduced the designed abundance gradient with a log– log ordinary-least-squares slope of k = 0.749, closer to the ideal slope of 1 than that obtained with pFind (k = 0.601). Both methods had a median absolute log2-fold error of 0.58, indicating similar overall absolute deviations, while π-MNovo better preserved the abundance dynamic range (Fig. 5a; Supplementary Method 10). Equal-input and abundance-gradient community profiles and replicate-averaged comparisons are shown in Supplementary Figs. S3, S4 and S5.

**Fig. 5.**
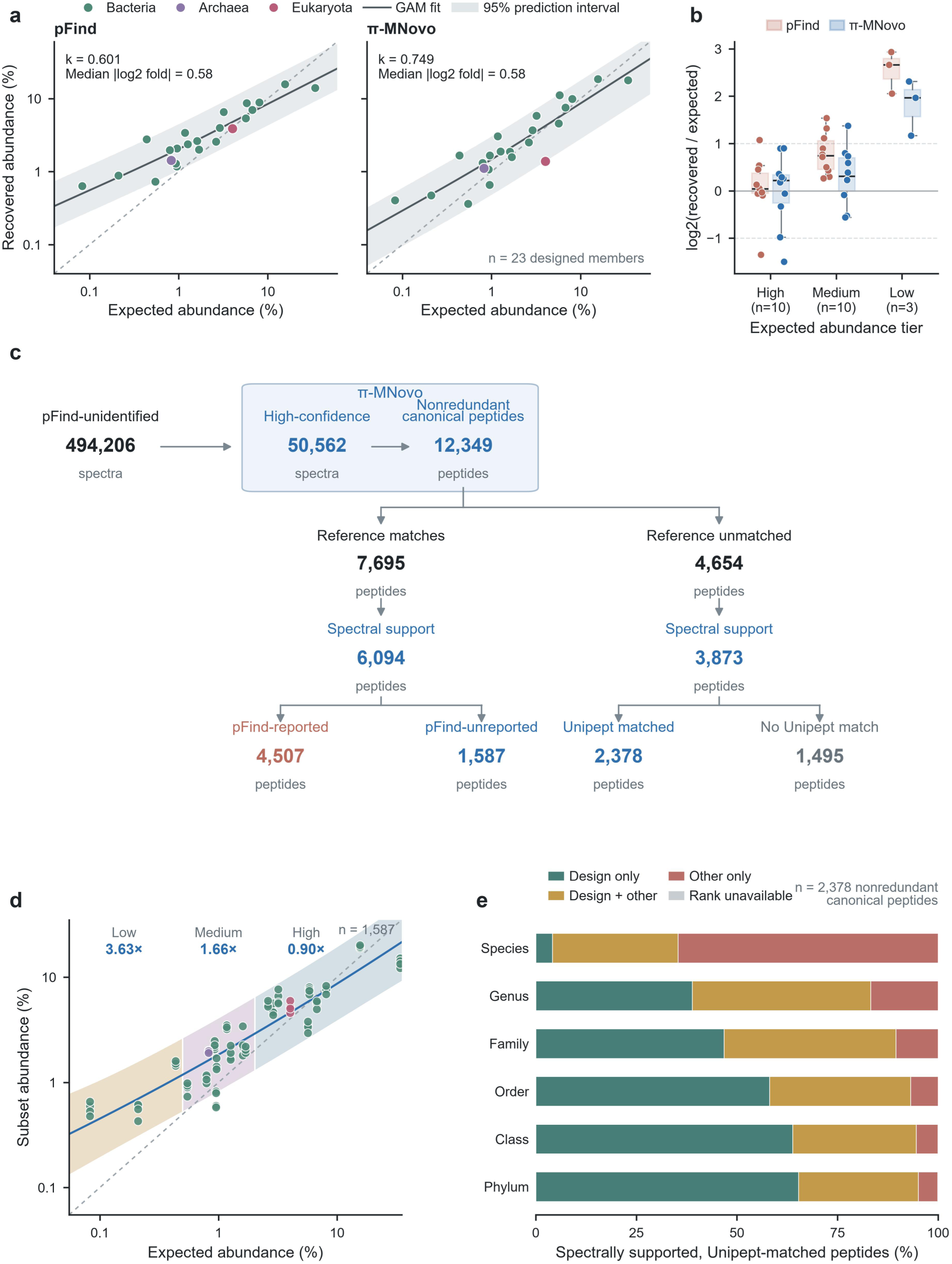
Synthetic-community abundance recovery and peptide evidence complementary to pFind. The bacteriophage-depleted reference contains 19 cellular members for P1–P4 and 23 for U1–U4. Panels a, b and d analyze U1–U4; c and e pool P and U communities. a, Expected and recovered abundances estimated separately for pFind and π-MNovo. Expected abundance denotes the designed species-level protein-abundance proportion reported for PXD006118. Points show member means across U1–U4 after averaging. Colors indicate Bacteria, Archaea and Eukaryota. Solid lines are generalized additive model (GAM) fits, shading shows pointwise 95% prediction intervals, and dashed lines indicate equality. k is the ordinary-least-squares slope from the regression of log10-transformed recovered abundance on log10-transformed expected abundance. Median |log2 fold| denotes the median absolute log2(recovered/expected) error. b, Recovery errors for pFind and π-MNovo, stratified by expected-abundance tier. Points represent individual members and show the median log2(recovered/expected) error across the eight U-series technical runs; boxes indicate the median and interquartile range, and whiskers extend to the most extreme values within 1.5 times the interquartile range. The solid line indicates zero error, and the dashed lines at −1 and 1 indicate twofold underestimation and overestimation. c, Evidence flow from pFind-unidentified spectra. High confidence denotes Score ≥ 0.92; canonical peptides are nonredundant I/L-equivalent sequences. Reference matches are exact matches to the cellular analysis reference. Spectral support requires at least one associated spectrum to pass the precursor and fragment criteria in Supplementary Method 11. pFind-reported and pFind-unreported denote peptides present in or absent from the accepted pFind peptide set across spectra, respectively. d, Quantification of 1,587 pFind-unreported, reference-matched peptides. Points show member abundances after averaging Run4 and Run5 within each community (92 observations). Point colors follow a; the line is a quadratic log–log fit. Shading shows pointwise 95% prediction intervals by abundance tier, and labels give median recovered/expected ratios. e, Rank-wise relationships between Unipept candidate taxa and the designed community for 2,378 spectrally supported, reference-unmatched peptides. Design only: all resolvable candidate taxa within the design; Design + other: candidates both within and outside; Other only: all outside; Rank unavailable: the rank cannot be resolved.

Recovery errors differed across the abundance range and between methods. For pFind, the median signed log2 recovery errors were 0.04, 0.74 and 2.66 in the high-, medium- and low-abundance groups, respectively; the corresponding values for π-MNovo were 0.22, 0.31 and 1.97. π-MNovo yielded median errors closer to zero in the medium- and low-abundance groups, whereas pFind was closer to zero in the high-abundance group (Fig. 5b). Both methods nevertheless overestimated the three low-abundance members, indicating that the larger errors at low abundance were not specific to de novo identification. Rather, these deviations may reflect cumulative biases across the measurement workflow, arising from differential recovery during sample preparation and abundance-dependent signal loss or undersampling during DDA acquisition. The low-abundance errors therefore cannot be attributed solely to π-MNovo.

We then examined sequence evidence recovered from pFind-unidentified spectra across P1–P4 and U1–U4 (Supplementary Method 11). π-MNovo produced high-confidence predictions for 50,562 of the 494,206 unidentified spectra, yielding 12,349 nonredundant canonical peptides. Of these, 7,695 matched the cellular reference database; 6,094 of the reference-matched peptides passed precursor-mass and fragment-evidence filtering. Of these 6,094 peptides, 4,507 were present in the accepted pFind peptide set, whereas 1,587 were not and were therefore classified as pFind-unreported peptides. Among the 4,654 peptides unmatched to the cellular reference, 3,873 passed the same spectrum-evidence filters; 2,378 had Unipept matches and 1,495 had none (Fig. 5c; Supplementary Table S5; Supplementary Data 8).

We next examined the abundance profiles obtained from the additional reference-matched peptides. When the 1,587 peptides not reported by pFind were quantified separately in U1–U4, the median recovered-to-expected abundance ratios were 3.63, 1.66 and 0.90 for low-, medium- and high-abundance members, respectively (Fig. 5d; Supplementary Fig. S6). Low-abundance members were overrepresented relative to the theoretical composition in this subset, consistent with the direction of the abundance errors in the full analysis. We separately examined the taxonomic affinities of the 2,378 Unipept-matched peptides that did not match the cellular reference. At the species level, 1,537 peptides (64.6%) were assigned exclusively to candidate taxa outside the designed community; overlap with designed taxa increased at broader taxonomic ranks (Fig. 5e; Supplementary Table S6). Among the spectrally supported, reference-unmatched peptides, 370 species-distinctive peptides mapped to 112 candidate species outside the designed community (Supplementary Data 9). Using the criterion of at least three distinctive peptides for taxonomic assignment21, *Paracoccus pantotrophus* (147 peptides) and *Staphylococcus epidermidis* (106 peptides) were each supported by more than 100 species-distinctive peptides, providing strong evidence that these taxa were present in the sample set but absent from the originally reported community. These analyses identified additional reference-matched peptides that contribute to community abundance profiles, together with reference-unmatched peptides that provided species-level taxonomic evidence beyond the experimental reference (Supplementary Method 12).

## Discussion

Across external microbial benchmarks and community analyses, π-MNovo improved peptide recovery by combining microbial-domain adaptation with evidence-guided selection from a shared candidate pool. This approach builds on advances in architecture, decoding and metaproteomic applications^4–8^ while addressing the practical task of choosing a complete sequence from several plausible predictions. The gains extended across species and peptide lengths and were accompanied by broader reference-based taxonomic coverage.

An important contribution of this study is the microbial spectral resource used for domain adaptation. Its design combines taxonomic breadth across 72 cultured gut bacterial isolates with broader peptide sampling within each isolate through five-fraction high-pH reversed-phase fractionation. Partitioning by canonical peptide sequence further prevents repeated spectra and modified forms of the same underlying peptide from occurring across development and internal test sets. Nevertheless, the resource represents cultivable human-gut isolates from five unevenly represented phyla and therefore does not capture the full diversity of microbial spectral space. Alongside the resource and the model, this study provides a metaproteomics-oriented evaluation framework. General de novo benchmarks such as NovoBench provide unified comparisons of sequence recovery, efficiency and robustness across proteomic datasets^22^. Our framework complements these evaluations by integrating tests on external microbial species, real metaproteomes and a synthetic community with known composition. Isolate spectra assess transfer across species and acquisition settings, whereas real metaproteomes assess performance in complex samples. The synthetic community provides a defined reference for interpreting recovered abundance profiles. These settings connect complete-peptide recall with reference-based taxonomic support and community reconstruction. Together with the microbial spectral resource, these evaluations provide a framework for assessing de novo models in metaproteomic applications.

Candidate availability and selection remain two areas for improvement. On the external benchmark, final recall was 66.30%, while the shared pool contained a reference-matching sequence for 68.13% of spectra. Better selection within the existing pool could therefore address the remaining 1.83-percentage-point gap. For the 31.87% of spectra without a reference-matching candidate in the pool, improvement would require generating such a candidate. The sequential component analysis identifies the gains associated with successive component additions.

Peptide length connects prediction difficulty with taxonomic information. Complete recovery became less frequent with increasing length, consistent with the challenge posed by incomplete fragment evidence^9^, whereas species-level reference assignments became more frequent. Together, these observations support efforts to improve long-peptide recovery in metaproteomic sequencing. The taxonomic information carried by a peptide also depends on homologous sequence sharing and reference-database composition^10^. Long peptides therefore represent a high-value but technically demanding regime: they are more difficult to recover completely, yet potentially provide more discriminating taxonomic evidence when correctly sequenced.

The synthetic community illustrates how de novo predictions can complement database searching^2–3^. π-MNovo recovered the overall abundance ranking with quantitative concordance comparable to that of pFind and contributed reference-matched peptides absent from the accepted pFind set. In this additional peptide subset, low-abundance members were overrepresented relative to the theoretical composition. The full analysis also showed larger abundance errors for low-abundance members. However, a similar pattern was observed for pFind, indicating that the larger deviations at low abundance were not specific to de novo identification. These deviations may reflect cumulative effects across sample preparation—including species-dependent protein recovery, digestion and peptide detectability—and DDA acquisition, particularly abundance-dependent precursor selection and undersampling. The added evidence therefore supports broader sequence coverage rather than improved quantitative accuracy. For peptides outside the cellular reference, overlap at higher taxonomic ranks suggests possible taxonomic affinities. Taxonomic matches alone do not establish the sequence correctness or organismal origin of these peptides.

The evaluation measures complete-system performance using released comparator weights and database-supported reference labels. Benchmark predictions were evaluated using a common residue-mass matching criterion. For external evaluations, backbone-training-unseen status was defined relative to the recorded backbone-training data. Score represents empirical sequence confidence, whereas the prediction bands describe model-conditional abundance relationships.

Together, the results support microbial-domain adaptation and evidence-guided candidate selection as a practical route to improving complete-peptide recall and expanding reference-supported evidence for metaproteomic interpretation.

## Methods

### Microbial spectral resource construction and reference annotation

The collection of human fecal samples was approved by the Ethics Committee of Beijing Institute of Technology (BIT-EC-H-2024004). Donors with acute or chronic gastrointestinal diseases, antibiotic use within the preceding three months, or other acute or chronic medical conditions were excluded. Written informed consent was obtained from all participants before sample collection. Fecal samples were collected following our previously established protocol for live microbiota biobanking^23^, and gut bacterial isolates were obtained from fecal samples collected from healthy adult volunteers.

We constructed the microbial spectral resource from cultured isolates and used it for microbial-domain adaptation, candidate-reranker training and internal testing. Briefly, biobanked fecal aliquots were recovered in eight different media under anaerobic conditions (5% H₂, 5% CO₂, and 90% N₂; 37 °C), serially diluted and plated on eight different agar media. Selected colonies were repeatedly re-streaked to obtain pure isolates, and a single colony from each isolate was expanded in the corresponding broth for downstream analyses. Cells were harvested by centrifugation at 16,000 × g for 20 min and washed three times with sterile PBS. The aliquoted pellets were stored at −80 °C until analysis. One pellet from each isolate was used for 16S rRNA gene sequencing of the V3 – V4 region using the universal bacterial primers 27F (5 ′ - AGAGTTTGATCCTGGCTCAG-3 ′ ) and 1492R (5 ′ -GGTTACCTTGTTACGACTT-3 ′ ), whereas a paired pellet was used for proteomic sample preparation. Thus, taxonomic identification and proteomic analysis were performed using paired aliquots derived from the same purified isolate culture.

Deep bacterial proteomic sample preparation and analysis are detailed in Supplementary Method 1. Briefly, after bacterial cell lysis, protein extraction, and digestion, peptides were separated into five fractions by high-pH reversed-phase C18 fractionation and analyzed by LC–MS/MS on an Orbitrap Exploris 480 in data-dependent acquisition mode. The raw data were analyzed using a two-step taxon-guided database-search strategy. In the first-pass search, MS/MS spectra were searched with pFind (3.2.2)^24^ against the MetaLab-MAG high-abundance protein (HAP) database derived from UHGG v2.0, which comprised ribosomal proteins and elongation factors. Identified peptides were subjected to taxonomic assignment using Unipept, with species-level assignments retained when sufficiently supported and genus-level assignments used otherwise. The Unipept assignments were cross-checked against the corresponding 16S rRNA gene sequencing results. Only isolates with concordant taxonomic assignments between the two approaches and with 16S rRNA gene sequencing results consistent with a pure culture were retained for subsequent database construction; isolates with discordant assignments or evidence of mixed cultures were excluded. For each retained isolate, the relevant UniProtKB^25^ proteome(s) corresponding to the consensus taxonomic assignment were retrieved and combined to construct a taxon-matched protein sequence database. The raw MS/MS data were then searched in a second pass with pFind against the resulting database. Peptide–spectrum matches were retained at an estimated PSM-level FDR of ≤ 1%.

Finally, the microbial spectral resource comprised 351 input MGF files from 72 bacterial isolates. Most isolates contributed five fraction-level files; 12 isolates contributed only three or four because 14 fraction samples were accidentally lost during experimental processing, whereas one isolate contributed ten files. During MGF annotation, we standardized peptide labels and representations of supported modifications, and converted I to L. We retained spectra with an absolute relative precursor-mass error of ≤ 100 ppm, calculated from the observed neutral mass and the sequence-derived neutral mass including supported modifications. The search tolerance was 20 ppm; the 100-ppm criterion was applied subsequently during annotation-file construction. Complete search and annotation settings are provided in Supplementary Table S1 and Supplementary Method 1.

### Data partitioning

We partitioned the data by canonical peptide key. Keys were defined after removing modifications, converting sequences to uppercase and treating I and L as equivalent. Replicate spectra and modified forms sharing the same key were kept in the same partition. A deterministic hash allocated peptide keys to training, validation and test sets with target proportions of 80%, 10% and 10%. After partitioning, we retained at most 120 spectra per key in the training set and at most five per key in each of the validation and test sets. The training, validation and test sets contained 9,144,192, 458,634 and 456,579 spectra, respectively. The corresponding numbers of unique canonical keys were 1,265,426, 158,998 and 158,401. No canonical key was shared among the three partitions. The final proportions of spectra reflect both the number of observations per peptide key and the split-specific caps.

Data within the training and validation partitions were assigned predefined roles in backbone adaptation, candidate reranking, router development and Score calibration. A four-project external development set was used for weight interpolation and selection of the high-confidence Score threshold. The 456,579-spectrum internal partition was excluded from component training and was used for frozen-component ablation and beam-width selection. Final comparative performance was evaluated on the independent external datasets. The four development projects are distinct from the external evaluation projects. The complete data-role map is provided in Supplementary Table S2 and Supplementary Method 2.

### Non-autoregressive Transformer backbone

We initialized π-MNovo from the publicly available π-PrimeNovo weights^7^ and retained its non-autoregressive Transformer encoder–decoder backbone. The encoder represents MS/MS peaks, whereas the decoder combines spectral representations, positional embeddings and precursor information to predict residue and blank-symbol probabilities in parallel. The decoder produces 40 CTC output positions. The database-search peptide-length range was 6–100 residues. A target with L model tokens and r adjacent identical-token pairs requires at least L + r CTC positions, so alignability in this decoder requires L + r ≤ 40. We trained the model with connectionist temporal classification (CTC)^26^, which sums over all generation paths that collapse to the reference peptide. Spectrum preprocessing and backbone settings are detailed in Supplementary Method 3.

CTC maps fixed-length positional outputs to variable-length peptide sequences. During path collapse, consecutive identical tokens are first merged and blank tokens are then removed, eliminating the need for residue-to-position alignment labels. The downstream rerankers use the same backbone outputs and candidate pool, separating peptide generation from candidate selection.

We adapted the backbone on microbial training spectra using a length curriculum that progressively increased the proportion of long peptides sampled. Complete-peptide recall on the validation set was used for model selection. After adaptation, we interpolated the base and fine-tuned weights as follows:

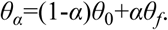

Here, θ_0_ and θ_f_ denote the base and fine-tuned weights, respectively. We evaluated the interpolation coefficient α on four external development projects and fixed it at 0.25. These projects were PXD034863^27^, PXD041951^28^, PXD011378^29^ and PXD019004^30^. We retained 5,000 spectra per project (20,000 in total), with one spectrum per canonical peptide key and no key overlap with the training, validation or internal test sets of the in-house resource. The same external development set was used to select the Score operating threshold. A fixed-budget development experiment informed the use of 1,000,000 spectra for backbone adaptation; further adaptation and interpolation details are provided in Supplementary Method 4.

### Candidate generation

π-MNovo generates peptide candidates through two decoding routes inherited from π-PrimeNovo: high-probability CTC beam search and precursor-mass-constrained (PMC) decoding^7^. The two outputs are merged and deduplicated to form a shared candidate pool for all downstream ranking paths.

Each of the three paths, R1, R2 and R3, rescores the shared pool and nominates a complete peptide sequence. This separation allows different evidence types to be compared using the same candidate pool.

### Evidence-guided candidate reranking

Each reranker leaves candidate sequences unchanged and learns only their relative order within the shared pool. A residual multilayer perceptron (MLP) predicts a candidate-specific correction to the baseline score for each path. The reranked score of candidate peptide y under path r is defined as:

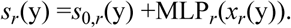

Here, r indexes the ranking path; *s_r_*(*y*) and *s*_0,*r*_(*y*) denote the reranked and baseline scores of candidate y, *x_r_*(*y*) is the path-specific feature vector and *MLP_r_* is the residual network for that path. R1 uses the length-normalized CTC score as its baseline. R2 and R3 use the R1 reranked score as their baseline and incorporate length-consistency and fragment-evidence features, respectively. Residual corrections were bounded and L2 regularization was used to limit departures from the baseline score. Rerankers were trained with a multi-positive listwise ranking loss:

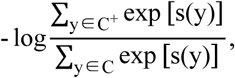

For a given spectrum, C is the valid candidate set and C⁺ contains candidates with the same length as the reference peptide and a residue-mass difference < 0.1 Da at every corresponding position. When more than one candidate meets this criterion, all are included in C⁺. The loss favors their combined probability over that of the remaining pool. Spectra without a candidate matching the reference peptide do not contribute to reranker optimization but remain in the denominator used for final evaluation.

The three paths share the residual reranking framework but use different types of evidence. R1 uses the length-normalized CTC score, the relative score difference from the top-ranked candidate, reciprocal rank and precursor-mass error. R2 adds a spectrum-conditioned peptide-length distribution and measures its agreement with candidate length. R3 matches theoretical b and y ions to observed peaks and summarizes the evidence using ion coverage, consecutive ion series and explained peak intensity. Reference peptides define the training labels and strata; inference uses only the input spectrum and candidate sequence. Candidate-pool construction and path-specific features are detailed in Supplementary Method 5.

### Conservative routing

To train the router, we compared the top-ranked predictions from R1, R2 and R3 with the reference peptide for each spectrum. Every path producing a complete reference match was treated as positive, and all positive paths formed the supervision set P. When no path matched, we used {R1} as the supervision set to encode the default-path fallback policy. The router was trained with a multi-positive path loss:

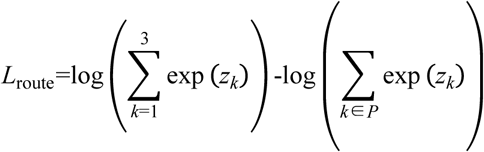

The three logits z are the unnormalized router outputs, and P is the supervised path set for the current spectrum. The loss maximizes the combined probability of the supervised paths. If all three paths are positive, this loss term is zero and provides no routing gradient. To balance the training categories, we retained every rescuable example in which R1 was incorrect and at least one expert was correct. Cases in which R1 was correct and at least one expert was incorrect were capped at twice the number of rescuable examples. Cases with three correct or three incorrect paths were capped at the number of rescuable examples.

A spectrum-level MLP with two hidden layers of 96 units each takes 36 input features derived from the spectrum, candidate sequences and three ranking paths. It produces three routing logits, which are normalized by softmax. R1 is the default path. The router switches to an expert only when the probability advantage of the highest-probability expert over R1 satisfies:

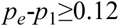

The expert probability is the larger of the R2 and R3 probabilities. Routing probabilities express relative preference among paths and are distinct from the final sequence Score. We selected δ using router-development data to maximize complete-peptide recall, subject to a harmful-switch rate of ≤ 0.1% across all spectra. Ties were resolved in favor of the lower switching rate. The released model uses δ = 0.12. Router architecture, supervision and data roles are detailed in Supplementary Method 6.

### Score calibration

We transformed the R1 sequence score associated with the final routed peptide into Score using a positive-slope Platt mapping fitted to 20,013 held-out spectra31. The mapping was fitted to R1 top-ranked candidates and then applied to the R1 score of the final selected candidate, regardless of which path selected it. After all components were frozen, we selected the operating threshold using final routed predictions on the separate four-project external development set of 20,000 spectra. The lowest threshold achieving at least 95% empirical complete-peptide precision was 0.917928; we rounded it up to 0.92 and used Score ≥ 0.92. This threshold was used only for high-confidence synthetic-community analyses; benchmark recall across eligible reference-annotated spectra was calculated without Score filtering. Score is a calibrated sequence-confidence value, distinct from an FDR or q value. Calibration and threshold selection are documented in Supplementary Fig. S7, Supplementary Table S7 and Supplementary Method 7.

### Benchmark evaluation and statistical analysis

All models were evaluated on the same eligible reference-annotated spectra using a common mass-aware matching rule. Residues were aligned from both peptide termini with cumulative mass differences < 0.5 Da and individual residue-mass differences < 0.1 Da; complete-peptide matching required equal sequence lengths and all residues to match. Complete-peptide recall is the number of matched predictions divided by the total number of evaluated spectra; invalid and empty predictions count as failures. At a given Score threshold, precision is the fraction of accepted predictions that match the reference, and coverage is the fraction of evaluated spectra with accepted predictions. Correct-peptide retention is the fraction of all correct predictions retained at that threshold. For the external seven-species benchmark, we treated species as fixed strata, resampled runs within species and then resampled spectra within the sampled runs. The same draws were used for all models. Two-sided 95% confidence intervals were obtained from the 2.5th and 97.5th percentiles of 5,000 bootstrap replicates (seed 3407). Bootstrap intervals were calculated separately for each model’s recall and for paired between-model recall differences. Figure 4b uses equal run weights and 10,000 run-bootstrap replicates across six fecal and 21 salivary runs; the resampling unit is the LC–MS/MS acquisition. Supplementary Table S3 reports run counts for the seven-species benchmark, and Supplementary Data 4 lists the 108 run identifiers and their evaluated spectrum counts. Further definitions are provided in Supplementary Method 8.

### Reference-based taxonomy and synthetic-community analysis

For the human-derived metaproteomes (PXD020786 and PXD055269), nonredundant reference-matched peptides were assigned to their lowest common ancestor (LCA) using Unipept 6.5.3^32–33^ and UniProtKB 2026.02. Rank counts are mutually exclusive rather than cumulative, and species support requires at least two species-LCA peptides.

In PXD006118, abundances were calculated separately from π-MNovo and pFind results. Shared-peptide PANDA intensities were allocated to species in proportion to the number of matching protein entries for each species, then normalized to sum to one over the designed cellular members within each run. Each biological-community replicate (U1–U4) had two technical acquisitions (Run4 and Run5). The technical replicates were averaged before calculating community-level summaries. The 1,587-peptide subset was quantified separately using the same allocation rule.

Peptides from pFind-unidentified spectra were deduplicated using I/L-equivalent canonical sequences. Reference matching used the cellular subset of the experimental reference after removal of bacteriophage proteomes. pFind-unreported peptides were defined as π-MNovo-predicted canonical peptides that matched the experimental reference database but were absent from the accepted pFind peptide set under the current search and filtering configuration. Both matched and unmatched branches were filtered using precursor error

≤ 20 ppm and empirical fragment thresholds. Within each peptide-length and precursor-charge stratum, up to 250 accepted pFind PSMs were selected by reservoir sampling with a fixed random seed. The strata, sampling procedure and empirical thresholds are specified in Supplementary Method 11 and Supplementary Table S8. The fifth percentiles of cleavage-site coverage and explained intensity were used as thresholds, and both criteria had to be met. For strata with fewer than 50 PSMs, thresholds were taken from the corresponding length-only group or, if that group was also too small, from the pooled reference PSMs. A peptide was retained if at least one associated spectrum met all criteria. Spectrum-evidence filtering assesses precursor and fragment consistency; an independent FDR was not estimated for the resulting de novo peptide set. Supplementary Methods 9–11 describe mapping, allocation, prediction-band construction and filtering.

## Data Availability

The isolate proteomic data generated in this study will be made publicly available after formal publication of this article. Public datasets analyzed in this study are available through ProteomeXchange under PXD037923, PXD012448, PXD014174, PXD028575 and PXD021084 (external microbial evaluation); PXD020786 and PXD055269 (human-derived metaproteomes); PXD006118 (synthetic community); and PXD034863, PXD041951, PXD011378 and PXD019004 (external development). Source Data for Fig. 5 provide panel-level plotting values, member-by-run abundance matrices and peptide-level summaries. Supplementary Data 1 provides training-resource statistics and partition summaries; Supplementary Data 2 provides 16S rRNA gene-based taxonomic identification and UniProtKB annotation; Supplementary Data 3 provides model configuration and checkpoint records; Supplementary Data 4 and 5 provide external run identifiers and benchmark statistics; Supplementary Data 6 provides ablation and routing results; Supplementary Data 7 provides real-metaproteome and taxonomy results; Supplementary Data 8 contains peptide- and spectrum-level evidence; and Supplementary Data 9 lists outside-design candidate species and their supporting peptide, spectrum and run counts. The files are listed in Supplementary Table S9.

## Code Availability

The complete π-MNovo model implementation and the trained model checkpoint used in this study are publicly available at https://github.com/PHOENIXcenter/pi-MNovo/tree/v0.1.0 (version v0.1.0; commit 636ca9c83c0b3cad46bd60b125ebb60808511fa6) under the MIT License. Scripts used for microbial-resource construction, canonical-peptide partitioning, benchmark evaluation and figure generation will be released in a versioned archive upon publication. Component training settings are summarized in Supplementary Table S10; checkpoint and source-record hashes are provided in Supplementary Data 3.

## Acknowledgements

This work was supported by the National Key Research and Development Program of China (2025YFA1309200 to L.L.), the Project Program of the State Key Laboratory of Medical Proteomics (SKLP-Y202401 to C.C. and SKLP-Y202402 to L.L.), and the Open Project of the National Center for Protein Sciences (Beijing) (2023-NCPSB-002 to L.L. and C.C.). The authors also acknowledge the technical support platforms at the National Center for Protein Sciences (Beijing), including the Mass Spectrometry Platform for assistance with mass spectrometry data acquisition, the Bioinformatics Platform for providing computational resources, and the Functional Proteomics Platform for assistance with microscopic observation.

## Author Contributions

J.Y., X.Z. and B.S. contributed equally to this work. J.Y. developed the π-MNovo framework, performed model training and computational analyses, curated the datasets, prepared the figures and drafted the manuscript. X.Z. performed bacterial isolation, purification and cultivation, 16S rRNA gene sequencing, proteomic sample preparation and mass spectrometry experiments, and drafted the sections describing the experimental procedures. B.S. contributed to the procurement and cultivation of bacterial isolates, preparation of fractionated proteomic samples, development of the quality-control pipeline and proteomic data analysis. T.L. provided technical guidance on π-PrimeNovo and advised on spectrum annotation, data preprocessing and model development. Z.L. advised on model construction and contributed key methodological ideas. J.D. curated the data for ProteomeXchange deposition and performed database searches of public datasets. Y.Z. and H.J. assisted with the wet-lab experiments. Z.H. contributed to the development of the microbial isolation protocol and standard operating procedures and was responsible for coordinating participant sample collection. L.J. co-supervised J.Y. as part of the joint training program. L.L. supervised the experimental work, and C.C. supervised the computational work. J.Y., L.L. and C.C. interpreted the results and revised the manuscript. All authors reviewed and approved the final manuscript.

## Competing Interests

The authors declare no competing interests.

## Notes

https://github.com/PHOENIXcenter/pi-MNovo/tree/v0.1.0

### Competing Interest Statement

The authors have declared no competing interest.

