## Supplementary Information for "π-MNovo improves de novo peptide sequencing through microbial-domain adaptation and evidence-guided candidate selection"

#### Supplementary Tables

##### Supplementary Table S1 | pFind search parameters used to construct the in-house microbial training resource

| Parameter | Setting |
|---|---|
| pFind version | 3.2.2 |
| Enzyme | Trypsin KR_C |
| Allowed missed cleavages | 3 |
| Precursor mass tolerance | 20 ppm |
| Fragment mass tolerance | 20 ppm |
| Fixed modification | Carbamidomethyl[C] |
| Variable modifications | Acetyl[ProteinN-term]; Oxidation[M] |
| Peptide-length range | 6–100 aa |
| Precursor-charge range | 2–5 |
| PSM-level FDR | 0.01 |
| Protein database | UniProt taxonomy-specific FASTA database |
| Contaminant and decoy handling | No contaminant sequences; standard reversed decoy database |

##### Supplementary Table S2 | Dataset roles and intended evaluation uses

| Role | Source | Spectra | Use | Evaluation role |
|---|---|---|---|---|
| Train | In-house microbial spectral resource | 9,144,192 | Parent set for model development | No |
| Validation | In-house microbial spectral resource | 458,634 | Epoch selection and Score calibration | No |
| Internal ablation set | In-house microbial spectral resource | 456,579 | Frozen-component ablation | Yes (internal) |
| Backbone adaptation | Fixed subset of the training role | 1,000,000 | Microbial-domain adaptation | No |
| Router | Training/selection/threshold roles | 116,784/49,913/49,914 | Training and selection of δ = 0.12 | No |
| Score calibration | R1 calibration subset | 20,013 | Platt mapping | No |

| Role | Source | Spectra | Use | Evaluation role |
|---|---|---|---|---|
| Score threshold | Four external development projects | 20,000 | Selection of 0.917928, applied as 0.92 | No |
| Interpolation development | Four projects: PXD034863, PXD041951, PXD011378 and PXD019004 | 20,000 | Selection of α = 0.25 | No |
| External benchmark | Seven species from five studies | 203,204 | Final external evaluation | Yes |
| Real metaproteomes | PXD020786 and PXD055269 | 734,769 | Complex-community evaluation | Yes |
| Synthetic community | PXD006118 | See source data | Community reconstruction and complementarity analyzes | Yes |

**Supplementary Table S3 | External seven-species benchmark and LC–MS/MS run structure**

| Project | Species | Strain | Instrument | Runs | Spectra |
|---|---|---|---|---|---|
| PXD012448 | Roseburia intestinalis | L1-82 | Q Exactive | 65 | 30000 |
| PXD014174 | Coprococcus eutactus | ART55/1 (deposited as Coprococcus sp. ART55/1) | Q Exactive Plus coupled to Ultimate 3000 RSLC nanoLC | 15 | 30000 |
| PXD021084 | Anaerostipes rhamnosivorans | DSM 26241 type strain | Q Exactive HF-X | 8 | 30000 |
| PXD028575 | Eubacterium maltosivorans | YI type strain (DSM 105863) | Orbitrap Exploris 480 coupled to EASY-nLC 1000 | 8 | 30000 |
| PXD037923 | Blautia hydrogenotrophica | DSM 10507 | Orbitrap Exploris 480 | 4 | 30000 |
| PXD037923 | Phocaeicola vulgatus | ATCC 8482 | Orbitrap Exploris 480 | 4 | 23204 |
| PXD037923 | Bacteroides uniformis | ATCC 8492 | Orbitrap Exploris 480 | 4 | 30000 |

Supplementary Table S3 summarizes project, species, strain, instrument and LC–MS/MS run counts for the analyzed external benchmark. The External_runs worksheet in Supplementary Data 4 lists all 108 Mascot generic format (MGF) TITLE-derived run identifiers, their species and projects, and the number of evaluated spectra per run. Reference annotation used the species-matched UniProtKB proteomes specified by the source annotation.

**Supplementary Table S4 | Overall recall on the external seven-species benchmark**

| Model | Spectra | Peptide recall (%) | 95% CI (%) |
|---|---|---|---|
| Casanovo | 203204 | 54.49 | 53.89–55.04 |
| InstaNovo+ v1.1.0 | 203204 | 56.43 | 55.78–57.05 |
| π-HelixNovo2 | 203204 | 58.55 | 57.92–59.13 |
| π-PrimeNovo | 203204 | 62.03 | 61.34–62.68 |
| π-MNovo | 203204 | 66.30 | 65.65–66.93 |

**Supplementary Table S5 | Evidence branches in the synthetic-community complementarity analysis**

| Evidence set | Operational criterion | Nonredundant peptides |
|---|---|---|
| High-confidence nonredundant π-MNovo peptides | All | 12,349 |
| Experimental-reference matches | Exact sequence match | 7,695 |
| Reference matched and spectrum-evidence filtered | Precursor and fragment evidence | 6,094 |
| Reported by pFind | Present in the qualified pFind results | 4,507 |
| Unreported by pFind | Absent from the qualified pFind results | 1,587 |
| Experimental-reference unmatched | No exact match in the bacteriophage-depleted experimental reference | 4,654 |
| Reference unmatched and spectrum-evidence filtered | Precursor and fragment evidence | 3,873 |
| Mapped by Unipept | At least one Unipept taxonomic assignment | 2,378 |
| No Unipept match | No Unipept taxonomic assignment | 1,495 |

**Supplementary Table S6 | Taxonomic consistency of peptides unmatched to the experimental reference but mapped by Unipept**

| Taxonomic rank | Design only | Design + other | Other only | Rank unavailable |
|---|---|---|---|---|
| Species | 98 | 743 | 1537 | 0 |
| Genus | 925 | 1055 | 398 | 0 |
| Family | 1112 | 1016 | 250 | 0 |
| Order | 1381 | 835 | 162 | 0 |
| Class | 1519 | 729 | 129 | 1 |
| Phylum | 1553 | 708 | 114 | 3 |

All percentages use 2,378 peptides that lacked an exact match to the cellular reference, passed spectrum-evidence filtering and received a Unipept assignment. Categories are assigned independently at each rank. Consequently, an outside-design species match can share a genus or higher-rank taxon with the designed community. Supplementary Data 9 lists outside-design candidate species, all matched peptide counts and species-distinctive peptide counts. Single-species matches require exactly one annotated candidate species in the recorded Unipept results, with no query cutoff or missing species annotations. Candidate matches are not used to confirm additional community members. Counts across species are not additive.

**Supplementary Table S7 | Selection of the Score operating threshold**

| Dataset | n | Use or criterion | Raw threshold | Precision | Correct-peptide retention |
|---|---|---|---|---|---|
| Four-project external development set | 20,000 | Lowest threshold with complete-peptide precision ≥ 95% | 0.917928 | 95.00% | 82.27% |

**Supplementary Table S8 | Empirical thresholds for spectrum-evidence filtering**

| Length (aa) | Charge | Reference PSMs | Cleavage coverage | Explained intensity |
|---|---|---|---|---|
| 06-10 | 2 | 250 | 0.666667 | 0.075072 |
| 06-10 | 3 | 250 | 0.666667 | 0.070463 |
| 11-15 | 2 | 250 | 0.636364 | 0.084401 |
| 11-15 | 3 | 250 | 0.571429 | 0.068073 |
| 11-15 | 4+ | 250 | 0.571429 | 0.071281 |
| 16-20 | 2 | 250 | 0.558681 | 0.095196 |
| 16-20 | 3 | 250 | 0.558681 | 0.065052 |
| 16-20 | 4+ | 250 | 0.485526 | 0.080998 |
| 21-25 | 2 | 250 | 0.300000 | 0.132985 |
| 21-25 | 3 | 250 | 0.464286 | 0.073933 |
| 21-25 | 4+ | 250 | 0.500000 | 0.073331 |
| 26+ | 2 | 250 | 0.200000 | 0.206165 |
| 26+ | 3 | 250 | 0.241379 | 0.087947 |
| 26+ | 4+ | 250 | 0.254891 | 0.072153 |

Thresholds are fifth percentiles expressed as fractions, not percentages. Values are displayed to six decimal places; full-precision values are supplied in Supplementary Data 8. Both fragment criteria and the precursor-mass criterion must pass. The listed reference strata each contained 250 usable PSMs; sparse strata use the fallback described in Supplementary Method 11.

**Supplementary Table S9 | Data files accompanying the manuscript**

| File | Contents |
|---|---|
| Supplementary Data 1 | Training-resource counts, partition summaries and internal data roles. |
| Supplementary Data 2 | 16S rRNA gene-based taxonomic identification and UniProtKB annotation of the 72 retained isolates. |
| Supplementary Data 3 | Release checkpoint, training configuration, Score calibration and source-record hashes. |
| Supplementary Data 4 | External datasets, comparator versions, 108 run identifiers and input-file hashes. |

| File | Contents |
|---|---|
| Supplementary Data 5 | Complete benchmark values, hierarchical bootstrap estimates and paired comparisons. |
| Supplementary Data 6 | Component ablations, candidate-width analysis, routing and candidate recovery. |
| Supplementary Data 7 | Real-metaproteome peptide performance and reference-based taxonomic evidence. |
| Source Data for Fig. 5 | Panel-level values, member-by-run quantification matrices and summaries for the 1,587- and 2,378-peptide subsets. |
| Supplementary Data 8 | Compressed peptide and spectrum evidence tables, intensity allocations and full Unipept results for Fig. 5. |
| Supplementary Data 9 | Outside-design candidate species, matched peptide counts and species-distinctive peptide evidence for Fig. 5e. |

Supplementary Data 1–7, Supplementary Data 9 and Source Data for Fig. 5 are Excel workbooks. Supplementary Data 8 is a ZIP archive containing tab-separated tables, gzip-compressed detail files and a field-level source manifest.

**Supplementary Table S10 | Component training configuration**

| Component | Learning rate | Batch | Epochs | Selected epoch | Seed |
|---|---|---|---|---|---|
| Backbone | $1.75 \times 10^{-4}$ | 96 | 5 | 5 | 20260730 |
| R1 | $3 \times 10^{-4}$ | 1024 | 8 | 5 | 20260731 |
| Length predictor | $3 \times 10^{-4}$ | 4096 | 8 | 8 | 20260731 |
| R2 | $3 \times 10^{-4}$ | 1024 | 19 | 19 | 20260731 |
| R3 | $3 \times 10^{-4}$ | 1024 | 60 | 38 | 20260731 |
| Router | $3 \times 10^{-4}$ | 1024 | 24 | 20 | 20260731 |

Epoch numbers are one-based. Epochs denotes the completed training schedule; selected epoch identifies the retained checkpoint. R2 and R3 training included at least 300 parameter updates. Rankers, the length predictor and the router used a learning rate of $3 \times 10^{-4}$ and weight decay of $10^{-4}$. The router used dropout 0.1. Detailed component settings and source-record hashes are provided in Supplementary Data 3.

### Supplementary Figures

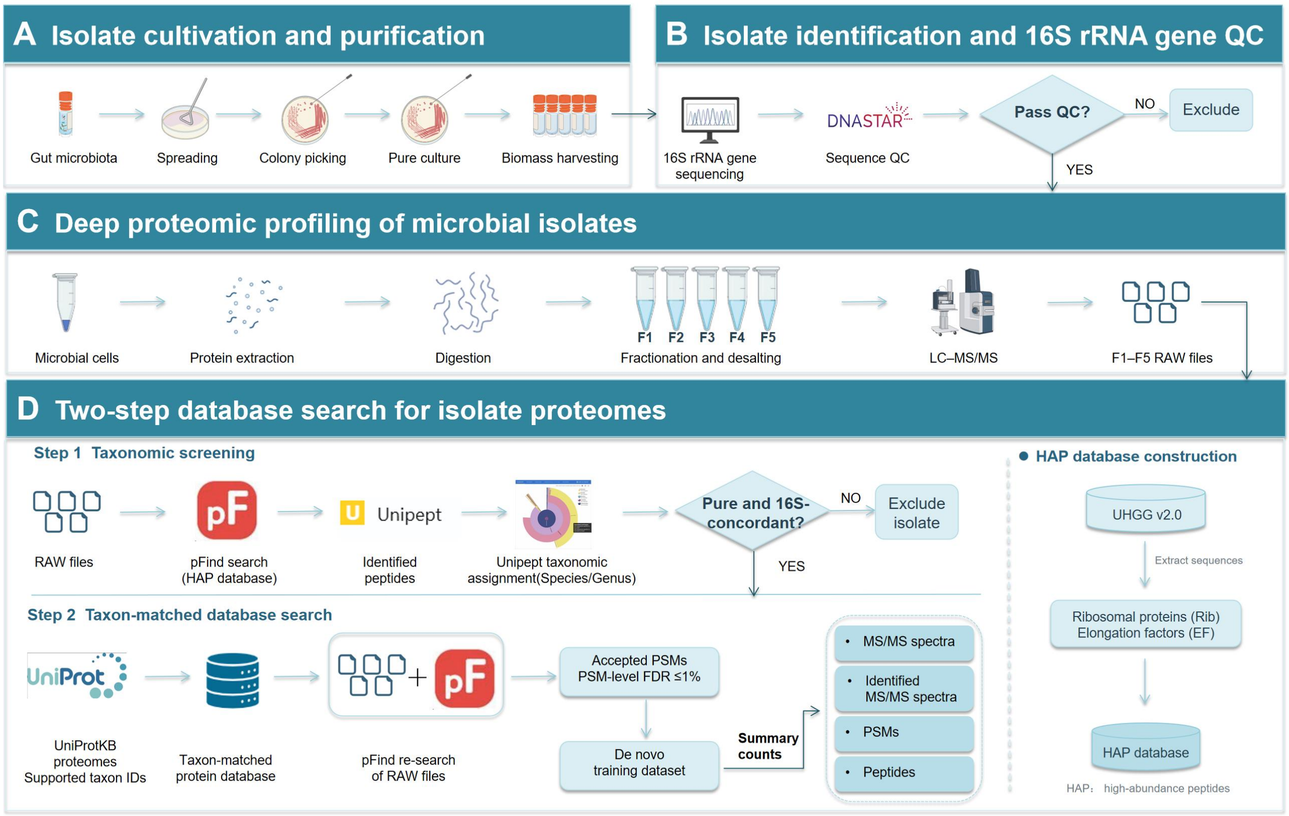


**Supplementary Figure S1 | Experimental workflow for constructing the peptide-annotated microbial MS/MS resource**

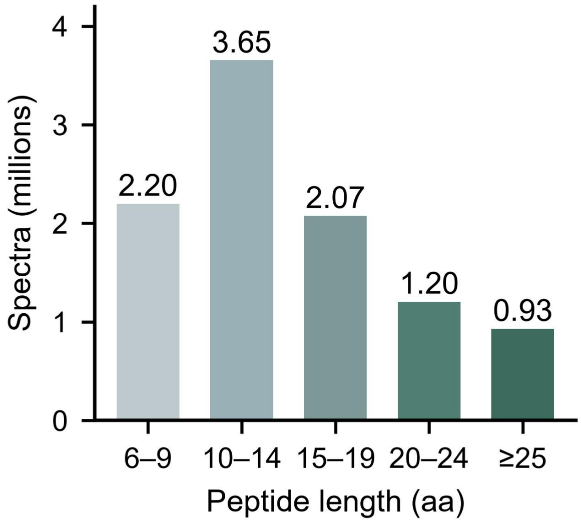


**Supplementary Figure S2 | Peptide-length distribution of the microbial spectral resource**

Distribution of the 10,059,405 retained spectra by canonical peptide length. Bar labels report millions of spectra.

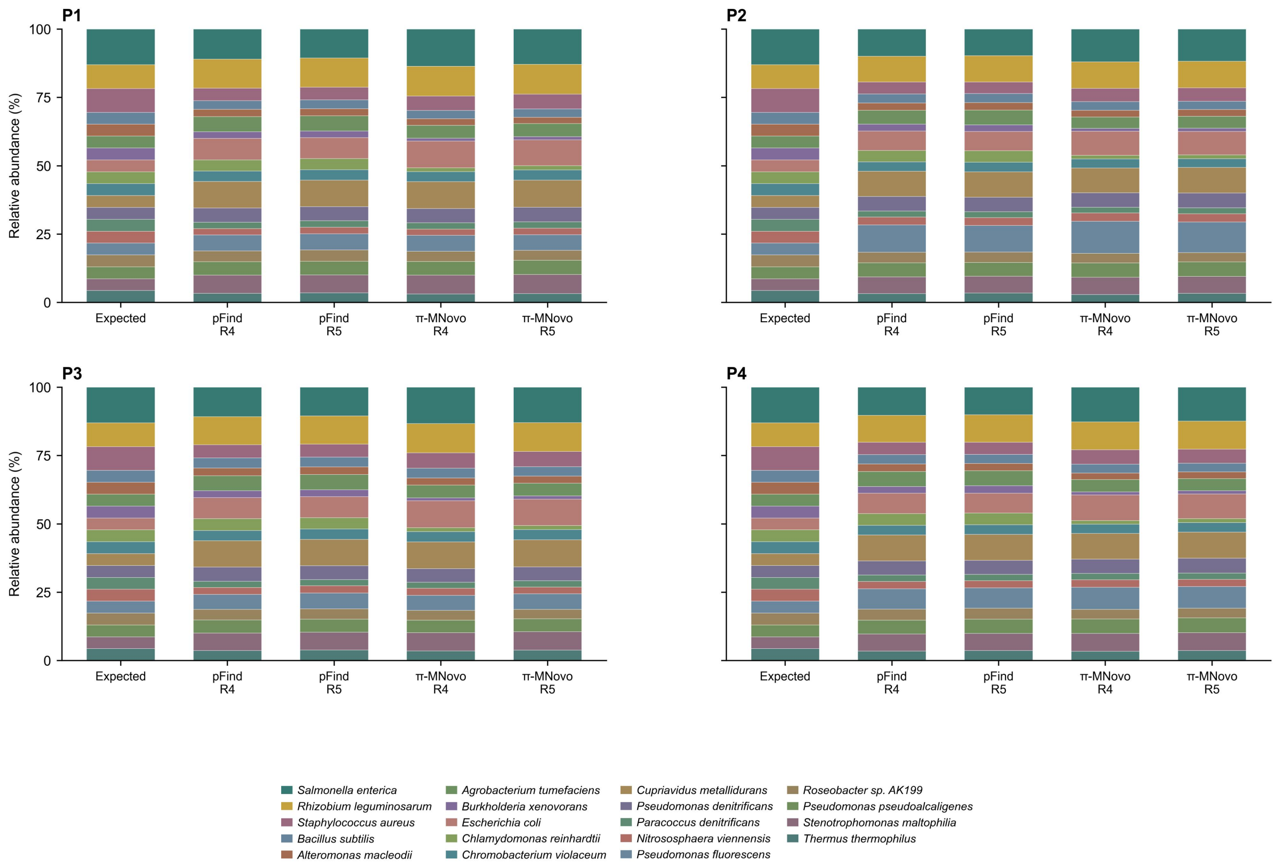


**Supplementary Figure S3 | Composition recovery in equal-input synthetic communities**

Cellular-member composition estimated separately by pFind and π-MNovo in the P1–P4 equal-input communities. Each bar represents a technical LC–MS/MS acquisition, and colors identify designed members. Relative abundance is closed over designed cellular members within each run. Run4 and Run5 are paired technical acquisitions from the same community sample.

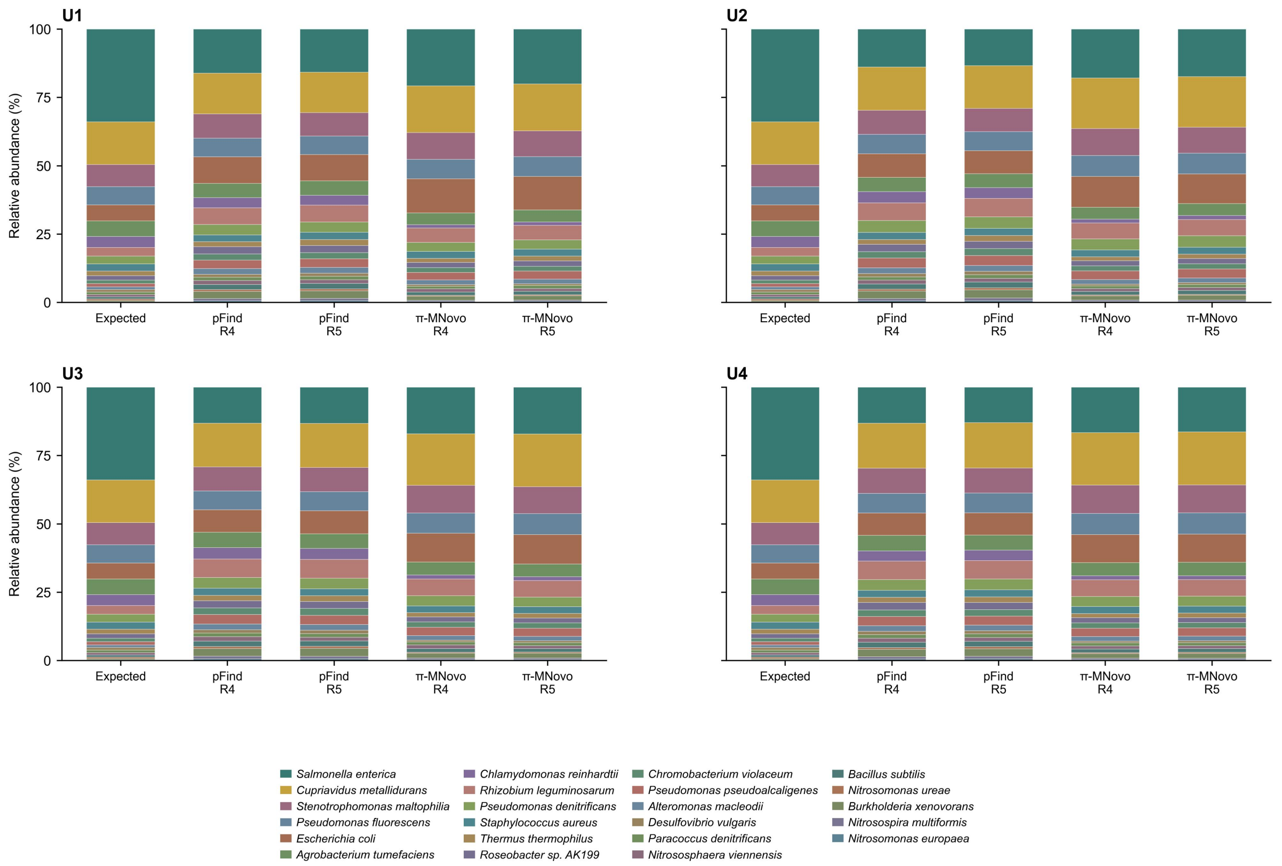


**Supplementary Figure S4 | Composition recovery in abundance-gradient synthetic communities**

Cellular-member composition recovered by pFind and π-MNovo in the U1–U4 abundance-gradient communities. Shared-peptide intensity was allocated in proportion to the number of matched protein entries per species and normalized over designed members within each run. Member colors are held constant across communities and methods. Run4 and Run5 are technical replicates.

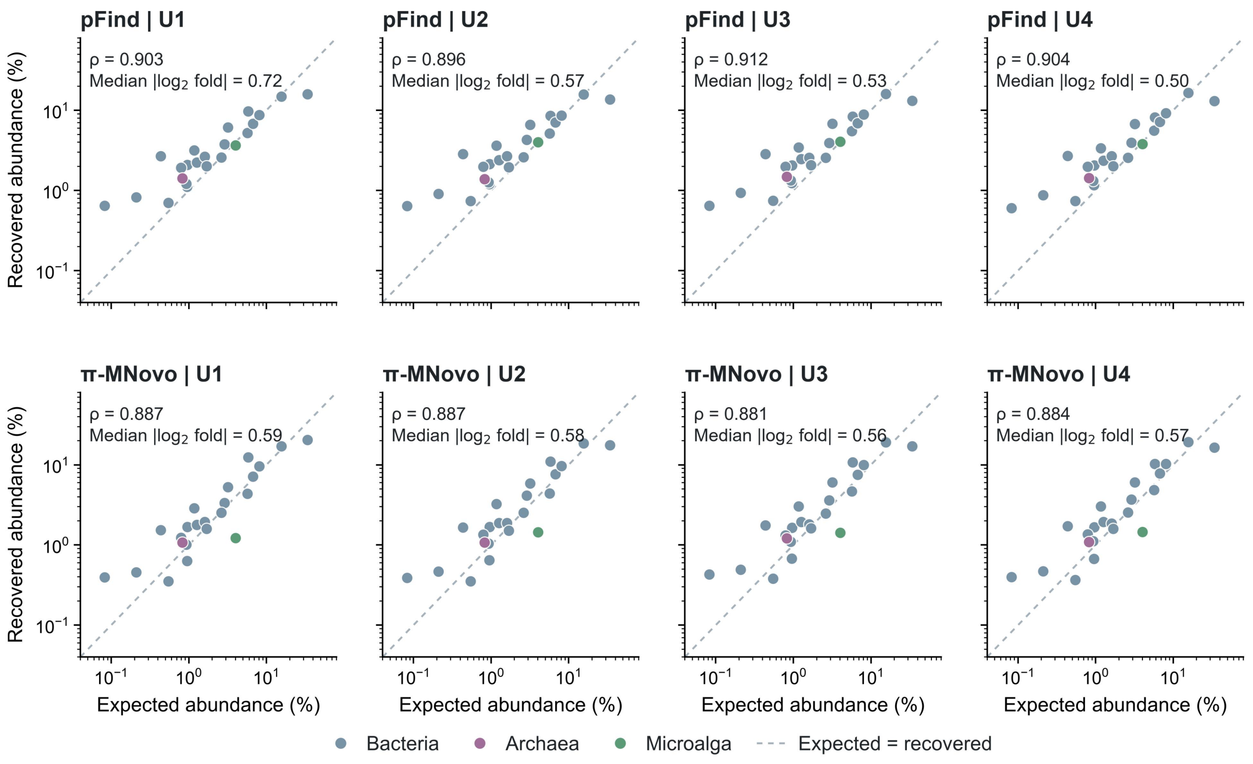


**Supplementary Figure S5 | Expected and recovered abundance after averaging technical replicates**

Within each U community and method, Run4 and Run5 abundances were averaged for each of the 23 designed cellular members. Colors distinguish bacteria, archaea and the microalgal member; the dashed line marks equality between expected and recovered abundances. Each panel reports Spearman ρ and the median absolute log2(recovered/expected) error across the 23 member-level means.

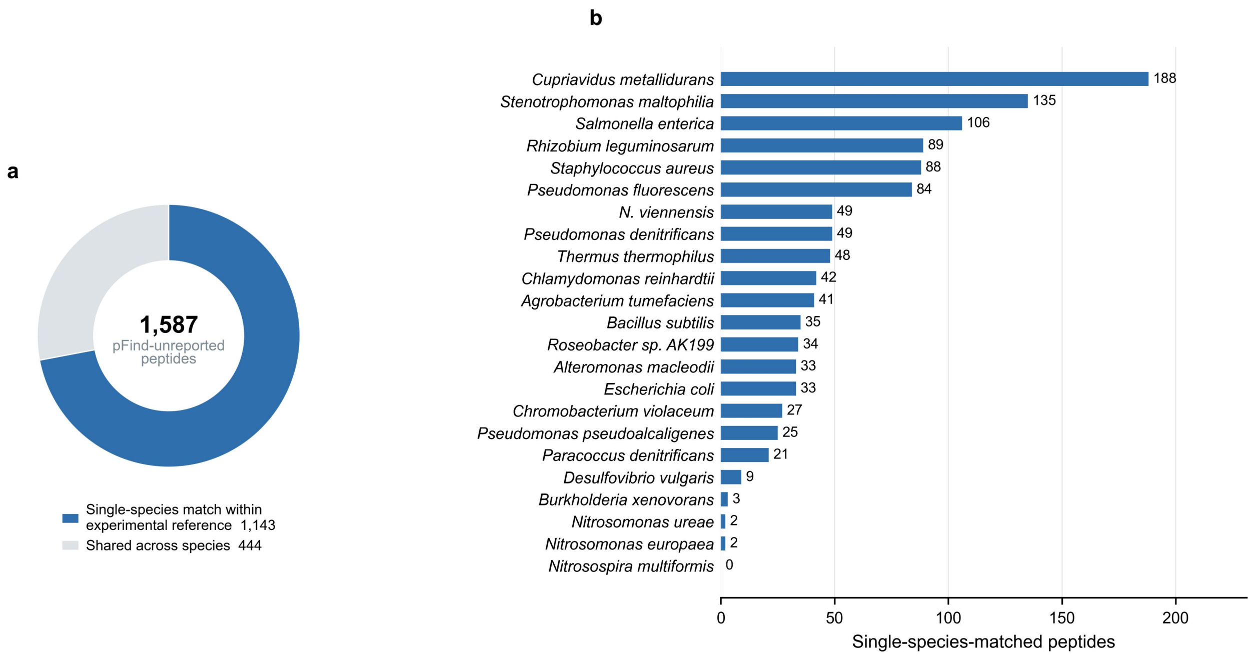


**Supplementary Figure S6 | Reference specificity of reference-matched peptides unreported by pFind**

a, Reference-mapping classes for the 1,587 peptides that matched the cellular reference, passed spectrum-evidence filtering and were absent from the accepted pFind peptide set. "Single-species match within experimental reference" denotes mapping exclusively to one species in the cellular analysis reference derived from Mock_Comm_RefDB_V3.fasta. b, Counts of these single-species-matched peptides by designed member. Shared peptides contribute to a only.

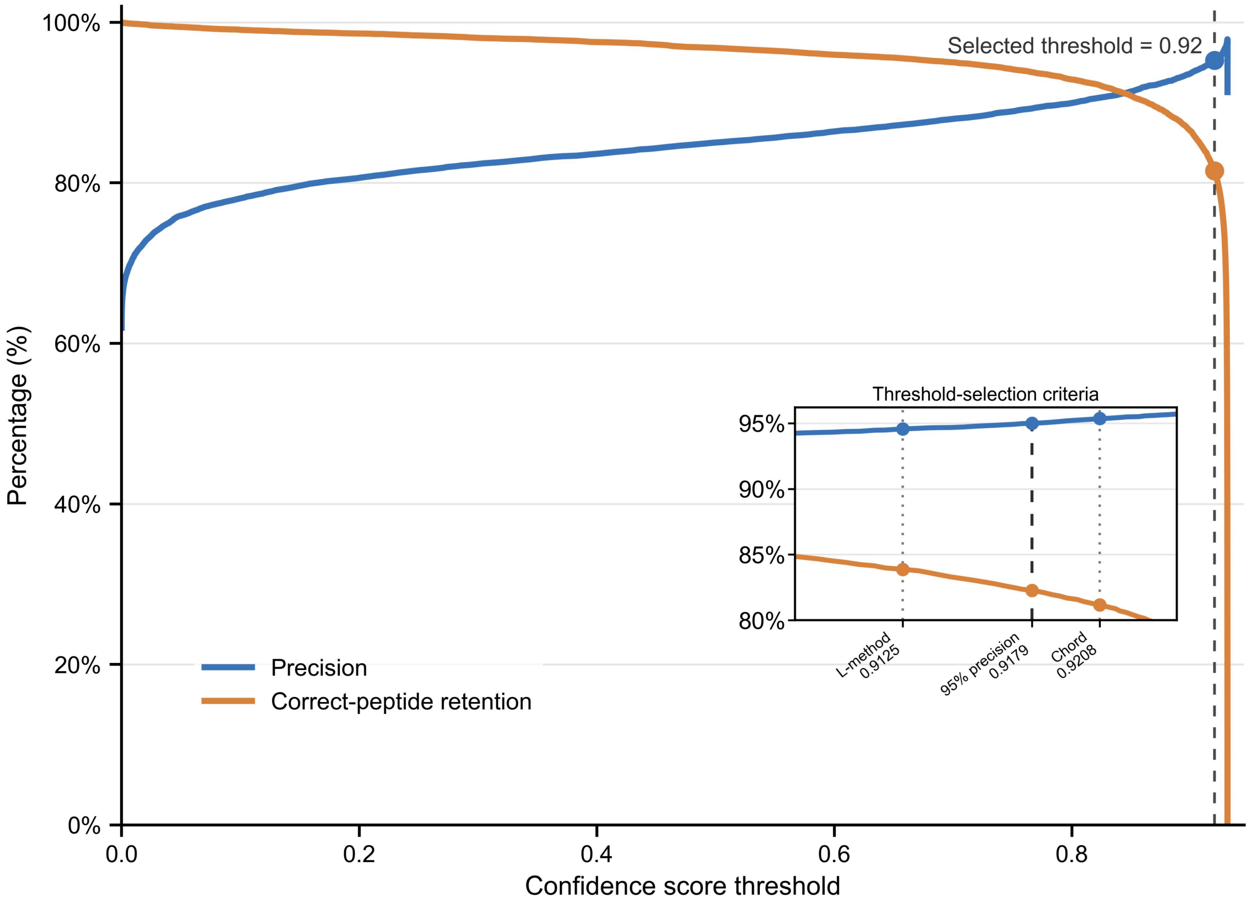


**Supplementary Figure S7 | Selection of the Score operating threshold**

Blue and orange curves show complete-peptide precision and correct-peptide retention for final routed predictions from 20,000 external development spectra, with 5,000 spectra from each of PXD034863, PXD041951, PXD011378 and PXD019004. All sequence-generation, ranking and routing components were frozen before threshold evaluation. Precision is the proportion of accepted predictions matching the reference; retention is the proportion of all correct final predictions retained above the threshold. The 95%-precision criterion gave a threshold of 0.917928 (0.9179 in the inset), with precision 95.0028%, retention 82.2697% and coverage 53.53%. The applied threshold was 0.92, shown by the main dashed line. The inset also displays L-method and chord-criterion thresholds as descriptive alternatives; the operating threshold was based on the precision criterion. Score is a calibrated sequence-confidence value; the percentages above describe the development-set operating criterion.

### Supplementary Methods

#### Supplementary Method 1 | Microbial sample preparation, LC–MS/MS analysis and spectrum annotation

##### Protein extraction and digestion

Proteomic samples were prepared using a previously reported metaproteomic workflow adapted for individual bacterial isolates[2]. Expanded bacterial cultures were centrifuged at 16,000 × g for 20 min at 4 °C, and the cells were washed twice with sterile PBS. After centrifugation at 300 × g for 5 min at 4 °C, the supernatants were transferred and centrifuged at 16,000 × g for 20 min at 4 °C to collect bacterial cells. The resulting pellets were stored at −80 °C overnight.

Cell pellets were lysed by sonication in 100 mM Tris-HCl (pH 8.0) containing 8 M urea, 4% (v/v) SDS and protease inhibitors. After centrifugation, proteins were precipitated overnight at −20 °C with five volumes of pre-cooled acetone/ethanol/acetic acid solution and quantified using a bicinchoninic acid (BCA) assay. For each sample, 100 µg of protein was reduced with 10 mM dithiothreitol (DTT), alkylated with 20 mM iodoacetamide (IAA) and digested with 4 µg of trypsin for 20 h at 37 °C. Peptides were desalted using C18 columns, eluted with 80% acetonitrile (ACN)/0.1% formic acid (FA) and dried under vacuum.

##### High-pH peptide fractionation

Desalted peptides were fractionated by high-pH reversed-phase C18 chromatography using published procedures[3,4]. Peptides were dissolved in 10 mM $NH_4HCO_3$ (pH 10) and loaded onto homemade C18 tips containing 7.5 mg of C18 material. Sequential elution used 6%, 9%, 12%, 15%, 18%, 21%, 25%, 30% and 35% ACN in 10 mM $NH_4HCO_3$ (pH 10). Fractions were combined as 6% + 21%, 9% + 25%, 12% + 30%, 15% + 35% and 18%, yielding five final fractions, which were dried under vacuum before LC–MS/MS analysis.

##### LC–MS/MS analysis

For each fraction, 1 µg of peptides was separated using a nanoEASY 1200 nano-flow liquid chromatography system coupled to an Orbitrap Exploris 480 mass spectrometer (Thermo Fisher Scientific) operating in data-dependent acquisition (DDA) mode. Mobile phases A and B were 0.1% FA in water and 0.1% FA in ACN, respectively. Peptides were separated using a 78-min gradient: 7% B at 0–6 min, 12–30% B at 6–57 min, 30–45% B at 57–67 min, 45–95% B at 67–68 min, 95% B at 68–75 min and 5% B at 75–78 min. MS1 spectra were acquired over m/z 350–1500 at a resolution of 60,000, with an automatic gain control (AGC) target of 300% and a maximum injection time of 50 ms. Precursor ions were fragmented by higher-energy collisional dissociation (HCD) with a normalized collision energy (NCE) of 27%[5].

##### Spectrum annotation and quality control

MS/MS spectra were searched with pFind against species-matched UniProtKB reference proteomes using the parameters listed in Supplementary Table S1. PSMs passing a 1% PSM-level FDR were retained. During generation of peptide-annotated spectra, the observed neutral precursor mass was recalculated from the precursor m/z and charge in the MGF file, and the theoretical mass was calculated from the reference peptide sequence and retained modifications. Records were retained only when the absolute relative neutral-mass error, calculated as $10^6$

× |m_obs − m_seq|/m_obs, was ≤ 100 ppm and both the sequence and modifications could be represented by the model vocabulary. This annotation-stage quality-control criterion was distinct from the 20-ppm precursor tolerance used for the pFind search.

**Supplementary Method 2 | Peptide-level partitioning and development roles**

A canonical peptide key was defined as the unmodified uppercase peptide sequence after treating isoleucine and leucine as mass-equivalent. Train, validation and test sets were first assigned deterministically at the peptide-key level so that no key occurred across splits. The three sets contained 9,144,192, 458,634 and 456,579 spectra, corresponding to 1,265,426, 158,998 and 158,401 unique peptide keys, respectively; pairwise cross-split overlap was zero. Training and validation data were subsequently assigned to backbone, ranker, expert, router and Score-development roles. Primary roles were peptide-key isolated wherever applicable, whereas secondary partitions for epoch selection, calibration or threshold selection followed the recorded split or key rule. Dataset roles and their evaluation status are summarized in Supplementary Table S2. The zero-overlap claim applies to the three primary partitions. External backbone-training-unseen status is defined separately by comparison against the backbone-training keys.

**Supplementary Method 3 | Spectrum preprocessing, vocabulary and backbone representation**

Each spectrum retained at most 800 peaks with nonnegative intensity over an m/z range of 1–6500. Peaks within ±1 Da of the precursor m/z were removed. Peak m/z values were encoded sinusoidally, and linearly projected intensities were fused with the peak representation. Precursor inputs comprised neutral mass, charge and m/z. The encoder and decoder each contained nine Transformer layers with a hidden dimension of 400, eight attention heads, a feed-forward dimension of 1,024 and dropout of 0.18. The parallel decoder emitted up to 40 positions. The vocabulary contained standard amino acids and the retained modified residues. Isoleucine and leucine remained separate model tokens but were treated as mass-equivalent for dataset partitioning and evaluation. The 40 decoder positions impose the CTC alignability condition $L + r \leq 40$, where L is the number of target model tokens and r counts adjacent identical-token pairs. Search parameters, including the annotation length range, are listed in Table S1.

**Supplementary Method 4 | Microbial-domain adaptation, length curriculum and weight interpolation**

Microbial-domain adaptation used a fixed subset of 1,000,000 spectra drawn in advance from the 7,799,398 spectra assigned to the backbone role. Pool membership remained fixed during adaptation. The fixed pool was sampled randomly without replacement from backbone_train only. Using zero-based epoch numbering, epoch 0 used uniform sampling with 36.62% long peptides; the sampling proportion was approximately 41.6% in epoch 1, 46.6% in epoch 2 and 50% from epoch 3 onward. Long peptides were defined as sequences of at least 16 residues. Weight interpolation was evaluated on 20,000 development spectra, comprising 5,000 spectra each from PXD034863, PXD041951, PXD011378 and PXD019004. Values of $\alpha$ = 0, 0.25, 0.50, 0.75 and 1.00 were compared, and $\alpha$ = 0.25 was fixed. These four projects were disjoint from the external seven-species benchmark, PXD020786, PXD055269 and PXD006118. A fixed-budget development experiment informed the use of the 1,000,000-spectrum subset for backbone adaptation. Adaptation used a batch size of 96, a learning rate of $1.75 \times 10^{-4}$, bfloat16 precision and seed 20260730. Training used eight data-loading workers and a maximum of five epochs. EarlyStopping monitored valid/pep_recall with min_delta=0.001 and patience=2. Both the best and last checkpoints were saved. The run comprised five epochs (52,085 steps), and the selected fine-tuned checkpoint was from epoch 5. Training settings are summarized in Supplementary Table S10 and Supplementary Data 3.

**Supplementary Method 5 | Mass-constrained candidate generation and ranking paths**

The union of Beam20 and precursor-mass-constrained candidates was used during training, whereas the union of Beam5 and precursor-mass-constrained candidates was used during inference. To select the inference beam width, beam widths of 1, 5, 10, 15 and 20 were compared on the internal ablation set with the backbone, R1 ranker and PMC configuration fixed. The corresponding peptide recalls were 65.801%, 65.881%, 65.900%, 65.912% and 65.920%, respectively. Beam5 was selected because increasing the beam width to 20 improved recall by only 0.0388 percentage points, corresponding to 177 additional correct predictions. Candidates were deduplicated by canonical peptide key, and all three ranking paths shared the same candidate pool and a single backbone forward pass. Positive training candidates had the same length as the reference peptide and a residue-mass difference < 0.1 Da at every corresponding position. The default path, R1, used the length-normalized CTC score, score difference from the original top-ranked sequence, candidate rank and precursor mass error. The length expert, R2, added the predicted length distribution and candidate-length consistency. The fragment-evidence expert, R3, evaluated theoretical b/y ions against experimental peaks using cleavage-site coverage, explained intensity, consecutive ion series and N- and C-terminal evidence. R2 and R3 reranked the shared candidates and reused the backbone predictions. The ranking networks used hidden width 96, dropout 0.05 and a bounded residual scale of 0.5, with a residual penalty coefficient of 0.001. The length predictor used hidden width 256 and dropout 0.1. Component-specific learning rates, batch sizes, epoch counts and seeds are listed in Supplementary Table S10.

**Supplementary Method 6 | Router supervision and conservative switching**

The router was a 36–96–96–3 multilayer perceptron. Inputs included top-ranked scores and score margins from the three paths, path agreement, precursor mass, charge, candidate length, predicted-length evidence, fragment evidence and precursor-mass-candidate source indicators. For each training spectrum, every path producing the correct peptide was treated as a positive routing label and optimized with a multi-positive cross-entropy objective. If all paths were incorrect, R1 was used as the fallback target. Samples for which R1 was incorrect but an expert was correct were retained, while the numbers of potentially harmful R1-correct expert cases and background cases were controlled to reduce class imbalance. The router used 116,784 spectra for training, 49,913 for epoch selection and 49,914 for selecting the switching margin $\delta = 0.12$. An expert replaced R1 only when its routing probability exceeded that of R1 by at least $\delta$. The margin was chosen to maximize peptide recall while constraining harmful switches to $\leq 0.001$ of all spectra. For all-correct examples, the supervised path set contains all three paths, and the displayed multi-positive loss contributes zero gradient. For all-incorrect examples, the R1 target encodes the fallback policy. The harmful-switch constraint was imposed on development data only.

**Supplementary Method 7 | Score calibration and operating threshold**

After fixing the rankers, a positive-slope Platt mapping was fitted to the raw R1 scores and correctness labels of R1 top-ranked candidates from 20,013 calibration spectra. Although the final prediction could be selected by R1, R2 or R3, its Score was always calculated by applying the same Platt mapping to that candidate's R1 score. Score did not affect candidate generation or routing. The operating threshold was selected after freezing all sequence-generation, ranking and routing components. Final routed predictions from 20,000 spectra, comprising 5,000 spectra from each of PXD034863, PXD041951, PXD011378 and PXD019004, were sorted by Score. The lowest threshold achieving complete-peptide precision ≥ 95% was 0.917928 (precision, 95.0028%; correct-peptide retention, 82.2697%; retained predictions, 53.53%) and was applied as Score ≥ 0.92 after rounding. These four projects were used for operating-threshold selection and were not part of the final external evaluation. The calibration fit and operating-threshold evaluation thus use separate data roles: R1 top-ranked candidates for fitting, and final routed predictions for threshold selection. Score is a calibrated sequence-confidence value, distinct from an FDR or q value.

#### Supplementary Method 8 | External evaluation and statistical analysis

The final external benchmark comprised 203,204 spectra from seven species, five published studies and four instrument configurations. All models received identical spectra and were evaluated with the same output parsing, I/L mass-equivalence rule and complete-peptide recall definition; empty or invalid predictions were counted as incorrect. Backbone-training-unseen denotes only that the corresponding canonical peptide key was absent from the π-MNovo backbone-training set; comparator pretraining overlap is outside this definition. Primary model differences were evaluated with 5,000 paired hierarchical bootstrap replicates and two-sided confidence intervals. Species were retained as fixed strata; LC–MS/MS runs were sampled with replacement within species, followed by sampling spectra with replacement within selected runs. All models shared each resampling index. The 95% confidence interval was defined by the 2.5th and 97.5th percentiles of the bootstrap distribution. The primary overall analysis was a spectrum-weighted micro-average. Paired recall differences and their percentile intervals form the primary statistical summaries. The 108 run identifiers and evaluated spectrum counts are provided in Supplementary Data 4, with species-level totals in Supplementary Table S3. Run identifiers were extracted from the TITLE fields of the evaluation MGF files using the same parser as the hierarchical bootstrap.

Mass-aware residue matching proceeded independently from the N and C termini. A residue match required a cumulative mass difference < 0.5 Da and an individual residue-mass difference < 0.1 Da. A prediction was counted as a correct complete peptide only when it had the same length as the reference and every residue matched. Amino-acid precision, amino-acid recall and complete-peptide recall were defined as follows.

$$\text{AA precision}=\frac{N_{\text{AA}}^{\text{correct}}}{N_{\text{AA}}^{\text{pred}}},\text{AA recall}=\frac{N_{\text{AA}}^{\text{correct}}}{N_{\text{AA}}^{\text{true}}}$$

$$\text{Peptide recall}=\frac{N_{\text{peptide}}^{\text{correct}}}{N_{\text{spectra}}^{\text{all}}}$$

The paired hierarchical bootstrap used random seed 3407. For Fig. 4b, the 27 LC–MS/MS runs were treated as equally weighted observations, and mean run-level recall differences and two-sided 95% percentile intervals were obtained from 10,000 nonparametric bootstrap resamples of runs using the same seed. Internal ablations were reported descriptively. Single-model recall intervals quantify uncertainty in recall, whereas paired difference intervals quantify between-model differences under shared resampling.

##### Supplementary Method 9 | Reference-based taxonomic annotation of real metaproteomes

Taxonomic mapping of peptides from real metaproteomes used Unipept 6.5.3 and UniProtKB release 2026.02, including reviewed and unreviewed entries. Unipept queries used extra=true, names=true, equate_il=true and tryptic=false. All valid peptide-to-protein and taxonomic mappings were collected for each canonical peptide, and reference-based taxonomic specificity was assigned using the lowest common ancestor. Each peptide was assigned to the most specific rank supported by all valid mappings. Assignments above phylum and peptides without a valid taxonomic mapping were grouped as other. Peptides were deduplicated within each project and again across projects when results were combined. A species-specific peptide had a unique species assignment within this reference-mapping space. A species was considered supported when at least two distinct correctly recovered peptides reached species-level LCA. The resulting counts measure reference-supported taxonomic coverage. The right-hand comparison in Fig. 4f counts species supported by π-MNovo but not by the corresponding comparator. Peptide-level mappings and taxonomic summaries are provided in Supplementary Data 7.

##### Supplementary Method 10 | Synthetic-community reconstruction and abundance processing

Synthetic-community analyzes used PXD006118. P1–P4 were equal-input communities and U1–U4 were abundance-gradient communities; Run4 and Run5 were technical replicates of the same sample. The cellular analysis reference was obtained by excluding bacteriophage proteomes. π-MNovo predictions were filtered at Score ≥ 0.92, and pFind results were filtered at 1% PSM-level FDR. Both used PANDA peptide intensities and exact I/L-equivalent mapping to the bacteriophage-depleted experimental reference derived from Mock_Comm_RefDB_V3.fasta. Member confirmation required at least two distinct species-LCA peptides; shared peptides contributed only to abundance estimation. Let Ip,s denote the raw PANDA intensity of peptide p in run s and np,k the number of protein entries in species k matched by peptide p. The allocation, species-level summation and within-run closure calculations are formalized below. The two methods were processed separately, using their respective PANDA peptide tables and within-method closure, rather than a π-MNovo–pFind union. After bacteriophage removal, P communities contain 19 cellular members and U communities contain 23. Figures 5a, b and d use U1–U4, whereas Figures 5c and e pool the P and U communities.

$$w_{p,k}=\frac{n_{p,k}}{\sum_j n_{p,j}}$$

$$I_{p,s,k}=I_{p,s}w_{p,k}$$

For a peptide matching only one species, the allocation reduces to:

$$I_{p,s,k}=I_{p,s}$$

$$Q_{s,k}=\sum_p I_{p,s,k}=\sum_p I_{p,s}w_{p,k}$$

$$A_{s,k}=\frac{Q_{s,k}}{\sum_j Q_{s,j}}$$

For U1–U4, relative abundance was calculated and closed within each run, then averaged over Run4 and Run5 for each member and community. For Fig. 5a, these values were further averaged across the four communities to obtain one expected/recovered pair per member. Spearman correlation and median absolute log2-fold error were calculated across the 23 members. Expected-abundance tiers were high (>2%), medium (0.5–2%) and low (<0.5%). A generalized additive model (GAM) was fitted in log10 space using a cubic regression spline with basis dimension 5 and restricted-maximum-likelihood smoothing. Pointwise 95% prediction bands used 5,000 bootstrap resamples of the four communities (seed 20260901), with smoothing fixed at the base-fit value. Each prediction draw combined a bootstrap fitted value with a sampled centered residual from the base GAM. The 2.5th and 97.5th percentiles of these draws define the band. For Fig. 5d, the same allocation and closure procedure was applied separately to the 1,587 pFind-unreported, reference-matched peptides. The 92 member–community combinations were fitted by quadratic ordinary least squares (OLS) regression in log10 space. Pointwise 95% prediction intervals used the Student t distribution with 89 degrees of freedom and incorporated residual variance and leverage. This model-conditional interval describes the observed abundance relationship. Source Data for Fig. 5 contain the panel-level points and fitted intervals, together with member-by-run allocated intensities and closed relative abundances.

##### Supplementary Method 11 | Database-search complementarity and spectrum evidence

Across the 16 P1–P4 and U1–U4 acquisitions, records were collapsed by raw-file identity and scan number so that multiple precursor-charge records from the same acquisition scan were counted once. This yielded 1,752,565 unique acquisition scans. A scan was classified as pFind-identified if it had at least one target PSM with q value ≤0.01; 1,258,359 scans met this criterion, leaving 494,206 pFind-unidentified scans for the complementarity analysis. Nonredundant π-MNovo and pFind peptide sets were constructed using the same canonicalization rule. A pFind-unreported peptide was absent from the accepted pFind peptide set across all analyzed spectra under the specified search configuration. Among 12,349 high-confidence nonredundant π-MNovo peptides, 7,695 exactly matched the experimental reference and 4,654 did not. Spectrum-evidence filtering required an absolute precursor-mass error ≤ 20 ppm and matching of theoretical b/y ions to experimental peaks within 20 ppm. Thresholds were derived from pFind reference PSMs accepted at 1% PSM-level FDR. The fifth percentiles of cleavage-site coverage and explained peak intensity defined the empirical lower bounds, and both criteria had to be satisfied. Peptide-length strata were 6–10, 11–15, 16–20, 21–25 and ≥26 residues; precursor-charge strata were 1, 2, 3 and ≥4. Accepted target pFind PSMs were restricted to supported modifications, eligible cellular-reference mappings and precursor-mass agreement within 20 ppm, and duplicate scans were removed. PSM files were read in lexicographic filename order and records in file order. Reservoir sampling retained at most 250 eligible PSMs per stratum using Python random.Random with seed 3407. After the first 250 records in a stratum, each subsequent record at position n replaced a uniformly selected reservoir entry with probability 250/n. The 3,500 usable reference PSMs occupied 14 strata with 250 PSMs each. Fifth percentiles were calculated by linear interpolation between adjacent ordered values at index $0.05(N-1)$. The final stratum-specific thresholds and reference counts are listed in Supplementary Table S8 and Supplementary Data 8. Strata with fewer than 50 PSMs used the corresponding length-specific thresholds, followed by global thresholds if the length stratum also contained fewer than 50 PSMs. Each peptide required at least one spectrum satisfying the precursor mass and both fragment-evidence criteria. Of the 7,695 reference matches, 6,094 passed these filters; 4,507 were reported by pFind and 1,587 were not. Among the 4,654 reference-unmatched peptides, 3,873 passed the same filters; 2,378 received a Unipept taxonomic assignment and 1,495 did not. Unipept queries used extra=true, names=true, equate_il=true and tryptic=false. Reference-unmatched status is defined against the cellular subset of the experimental reference. Peptide-level summaries and rank-specific assignment categories are provided in Source Data for Fig. 5; the associated spectrum evidence, intensity allocations and full Unipept results are provided in Supplementary Data 8.

##### Supplementary Method 12 | Protein evidence and implementation

The 1,587 reference-matched, spectrum-supported and pFind-unreported peptides were mapped by exact sequence to reference proteins and classified as unique-protein or multi-protein-compatible mappings. These mappings describe peptide support for compatible reference protein entries. Models were implemented with PyTorch and PyTorch Lightning. The unified v0.1.0 checkpoint contains the backbone, rankers, length predictor, router, normalization parameters, vocabulary and inference settings. Supplementary Data 3 provides its SHA256 digest and the training-summary hashes for the component records.
